# Engineering self-propelled tumor-infiltrating CAR T cells using synthetic velocity receptors

**DOI:** 10.1101/2023.12.13.571595

**Authors:** Adrian C. Johnston, Gretchen M. Alicea, Cameron C. Lee, Payal V. Patel, Eban A. Hanna, Eduarda Vaz, André Forjaz, Zeqi Wan, Praful R. Nair, Yeongseo Lim, Tina Chen, Wenxuan Du, Dongjoo Kim, Tushar D. Nichakawade, Vito W. Rebecca, Challice L. Bonifant, Rong Fan, Ashley L. Kiemen, Pei-Hsun Wu, Denis Wirtz

## Abstract

Chimeric antigen receptor (CAR) T cells express antigen-specific synthetic receptors, which upon binding to cancer cells, elicit T cell anti-tumor responses. CAR T cell therapy has enjoyed success in the clinic for hematological cancer indications, giving rise to decade-long remissions in some cases. However, CAR T therapy for patients with solid tumors has not seen similar success. Solid tumors constitute 90% of adult human cancers, representing an enormous unmet clinical need. Current approaches do not solve the central problem of limited ability of therapeutic cells to migrate through the stromal matrix. We discover that T cells at low and high density display low- and high-migration phenotypes, respectively. The highly migratory phenotype is mediated by a paracrine pathway from a group of self-produced cytokines that include IL5, TNFα, IFNγ, and IL8. We exploit this finding to “lock-in” a highly migratory phenotype by developing and expressing receptors, which we call velocity receptors (VRs). VRs target these cytokines and signal through these cytokines’ cognate receptors to increase T cell motility and infiltrate lung, ovarian, and pancreatic tumors in large numbers and at doses for which control CAR T cells remain confined to the tumor periphery. In contrast to CAR therapy alone, VR-CAR T cells significantly attenuate tumor growth and extend overall survival. This work suggests that approaches to the design of immune cell receptors that focus on migration signaling will help current and future CAR cellular therapies to infiltrate deep into solid tumors.

## INTRODUCTION

Chimeric antigen receptor (CAR) T cells express antigen-specific synthetic receptors, which upon binding to cancer cells, elicit T cell antitumor responses^1^. CAR T cell therapy has enjoyed success in the clinic for hematological cancer indications^1^, giving rise to decade-long remissions in some cases^2^. However, CAR T therapy for patients with solid tumors has not seen similar successes^1,3^. Solid tumors constitute 90% of adult human cancers^4^, representing an enormous unmet clinical need ripe for new cell therapies.

Many recognize that infiltration of immune cells into the tumor may be key for effective anti-tumor activity^5,6^. Since CAR T cells require direct contact with target tumor cells for subsequent killing, we hypothesize that the discrepancy in success of CAR T therapy between liquid and solid tumors is related to the CAR T cell’s ability to physically enter and penetrate deeply into the tumor microenvironment (TME) (Fig. 1a). In solid tumors, the TME contains a highly dense and complex extracellular matrix (ECM), the major non-cellular constituent of the TME^7,8^, which behaves as a physical barrier to T cell migration^9^ (Fig. 1a compares the physical barriers in the TMEs of liquid tumors and solid tumors). This tumor-associated collagen-rich stromal matrix is generally denser and stiffer than most normal tissues^10^, and T cells demonstrate decreased motility^11^ and infiltration^12^ within stiff collagen matrices^13–15^.

**Fig. 1.**
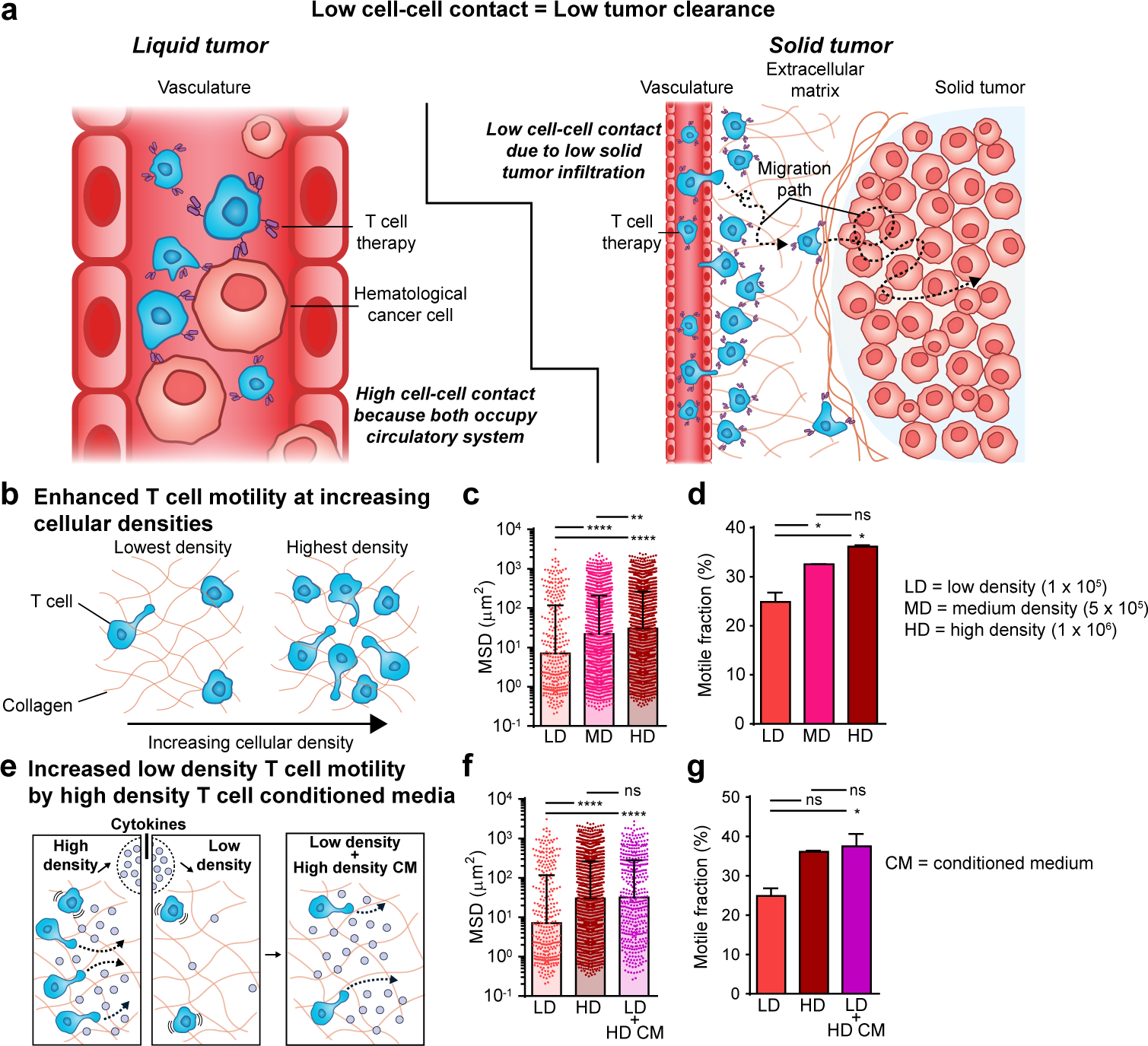
T cells migrate in a density-dependent manner *in vitro*. **a**, Current T cell therapies need to traverse a major physical barrier – the extracellular matrix (ECM) – in the tumor microenvironment (TME) of solid tumors for eventual cell-cell contact that they do not experience in the context of liquid tumors. **b,** Primary human pan T cells were isolated from primary human PBMCs using a donor leukopak. Isolated T cells premixed with a CD2/CD3/CD28 activation solution and 100 IU/ml IL2 were encapsulated in 3D gels composed of the major ECM protein, type 1 collagen (rat tail type 1 collagen at 2 mg/ml), at increasing T cell densities. **c,** After a 48-h incubation, the spontaneous motion of T cells at increasing densities (LD = 10^5^ cells/ml, MD = 5×10^5^ cells/ml, HD = 10^6^ cells/ml) embedded in 3D collagen gels was monitored using time-lapsed phase-contrast microcopy for 1 h in 3-min intervals at 37 °C and 5% CO_2_. Cells were manually tracked to generate trajectories for each cell (see Extended Data Fig. 1). A custom software was used to convert trajectories into mean squared displacement (MSD) for each cell at a time lag of 9 min. Each dot represents an individual cell. LD = low density, MD = medium density, HD = high density. **d,** A Matlab script was coded to extract the fractions of T cells moving more than their own size (>R^2^ = 25 μm^2^, where R = 5 μm is the average T cell radius) from the MSDs shown in panel (**c**) (see also Extended Data Fig. 1). **e,** Activated LD and HD T cells in 3D collagen gels were incubated for 48 h, after which medium from LD T cells was replaced with conditioned medium (CM) obtained from HD cells. **f,** MSDs of LD, HD, and LD + HD CM T cells imaged and analyzed (as in **c**). **g,** Motile fractions corresponding to panel (**f**), computed as in panel (**d**). For all experiments MSD, median with the SEM are plotted (*n*=2 technical replicates per biological replicate, *N*=2 biological replicates). Individual dots represent individual cells; an average of at least 150 cells per technical replicate, corresponding to up to 989 cells per technical replicate were tracked (see source data). One-way ANOVA with Dunn’s multiple comparison test was used for statistical analysis (ns = not significant, *\*P* < 0.01, *\*\*P* < 0.01, *\*\*\*\*P* < 0.0001). For all experiments measuring the motile fraction, mean with SEM is plotted (*n*=2 technical replicates per biological replicate, *N*=2 biological replicates).

A recent effort to enhance tumor infiltration and accumulation involved engineering CAR T to express synthetic notch receptors that induce local T cell proliferation via IL2 secretion in the TME^16^. Another effort aimed to target and deplete cancer-associated fibroblasts in the stroma that surrounds the tumor^17^. However, these approaches do not solve the central problem of limited ability of therapeutic cells to migrate through the stromal matrix^3^. A step towards solving this problem is the strategy of enhanced chemotaxis of therapeutic cells toward the tumor site^18,19^, but chemotaxis and migration, while often confused with one another, are not interchangeable^20^. Chemotaxis refers to biased, directional migration along chemical gradients, while migration is unbiased (random, nondirectional) movements that occur in the absence of chemical gradients^20^. A cell that can chemotax towards cancer cells but has limited or no migratory capability is akin to steering a car’s wheel with its engine turned off or a car in low gear.

Here we address directly the challenge of tumor infiltration by engineering CAR T cells to have a fast migration phenotype. First, we discovered that T cells at low and high density display low- and high-migration phenotypes, respectively. The highly migratory phenotype is mediated by a paracrine pathway from a group of self-produced cytokines that include IL5, TNFα, IFNγ, and IL8. Then, we exploit this finding to “lock-in” a highly migratory phenotype by developing and expressing receptors, which we call velocity receptors (VRs). VRs target these cytokines and signal through these cytokines’ cognate receptors to increase T cell motility by increasing the fraction of cells into the high-migration phenotype. In preclinical mouse models, we establish that VR-CAR T cells can infiltrate lung, ovarian, and pancreatic tumors in large numbers and at doses for which control CAR T cells remain largely confined to the tumor periphery. In contrast to CAR therapy alone, VR-CAR T cells significantly attenuate tumor growth and extend overall survival. This work suggests that approaches to the design of immune cell receptors that focus on migration signaling will help current and future CAR cellular therapies to infiltrate deep into solid tumors.

## RESULTS

### More T cells migrate at high density

Before we could develop a highly motile CAR T cell, we first needed to understand how normal T cells migrate in 3D collagen-rich settings *in vitro*. We hypothesized that identifying basic biophysical principles of enhanced T cell migration in simple *in vitro* models could help us design new CAR T cells featuring much higher infiltration potential *in vivo*. Using time-resolved phase contrast microscopy, we tracked the spontaneous movements (no added chemokines or chemo-attractants) of hundreds of unlabeled primary human T cells purified from healthy donors, activated by CD3/CD28 beads, and fully embedded in 3D collagen matrices (see details in Materials and Methods). This standard 3D collagen matrix model has been used extensively for cell migration studies^21–24^ and serves here as a screening platform. The mean squared displacement (MSD) of each cell was computed from the trajectories of cells in the matrix (Extended Data Fig. 1a) and used as the primary parameter to compare the motility of individual cells. Due to the non-Gaussian nature of the distribution of cellular movements, traditional migration parameters such as speed and persistence time/distance could not be extracted. T cells exhibited heterogeneous migratory patterns^25^, where some T cells migrated more than their own size, ∼5 μm, during recording time, while others hardly moved at all, suggesting different migratory states.

Recent work has shown that cell density (number of single cells in a set volume) can modulate the migration of cancer cells in collagen matrices^26^. Breast cancer cells and fibrosarcoma cells showed a small, ∼1.5-fold increase in cell speed for a 10-fold increase in cell density^26^. Here, we explored the possibility that cell density could modulate the migration of T cells (Fig. 1b). We found that T cells increased their migration up to 4-fold, as cell density was increased 10-fold, from 10^5^ to 10^6^ cells/ml (Fig. 1c and Extended Data Fig. 1a,b). Though not necessarily at physiological levels, we chose these densities to elicit a potential change in cell speed that would help us in the downstream identification of molecules responsible for this change. Furthermore, these cell densities were chosen to provide enough space between cells such that direct cell-cell collisions were rare. We observed that the MSD of T cells that migrated more than the square of their radius did so similarly across all tested densities (Extended Data Fig. 1c,d). This suggested that T cell migration adopted two main states: a high-motility state corresponding to the same high MSD (> R^2^, where R ∼ 5μm is the T cell radius) or a low-motility state for which significantly fewer cells were motile (MSD < R^2^). Thus, the overall difference in population-averaged cell MSD at different densities depended on the percentage of cells in a high-motility state (Fig. 1d).

Since we observed limited T cell intercellular collisions, we hypothesized that this density-induced increase in cell migration was caused by secreted molecules. To test this hypothesis, we cultured T cells at low density (LD) in a collagen matrix with conditioned medium (CM) harvested from T cells cultured at high density (HD) in a collagen matrix (Fig. 1e). This conditioned medium (CM) was sufficient to increase the motility of T cells at LD to levels similar to T cells at HD (Fig. 1f and Extended Data Fig. 1e,f). Once again, this increase in population-averaged MSD corresponded to a shift in the number of cells in the high-motility state (Fig. 1g and Extended Data Fig. 1g,h) This result suggested that factors secreted by cells at HD increased the percentage of cells in a high motility state.

### High T cell migration can be induced by pro-inflammatory cytokines

Next, we sought to identify the factors secreted by cells at HD responsible for their increased motility. We utilized a microchip to run single-cell proteomics^27,28^ on the medium from LD cells and HD cells suspended in 3D collagen matrices (Fig. 2a). We identified four cytokines – IL8, TNFα, IFNγ, and IL5 – that were secreted by HD cells but undetectable for LD cells (Fig. 2b), suggesting their *de novo* production by T cells at a high cellular density. The protease granzyme B, which remodels the ECM, thereby allowing more rapid T cell motility^29^, was also detected for HD cells and undetected for LD cells. But, given our aim to engineer receptors that induce T cell migration signaling, we decided to not pursue granzyme B as it has no known cognate receptor. IL8 is a well-established T cell chemoattractant^30^, but little is known about its effect on T cell migration (i.e. not chemotaxis).

**Fig. 2.**
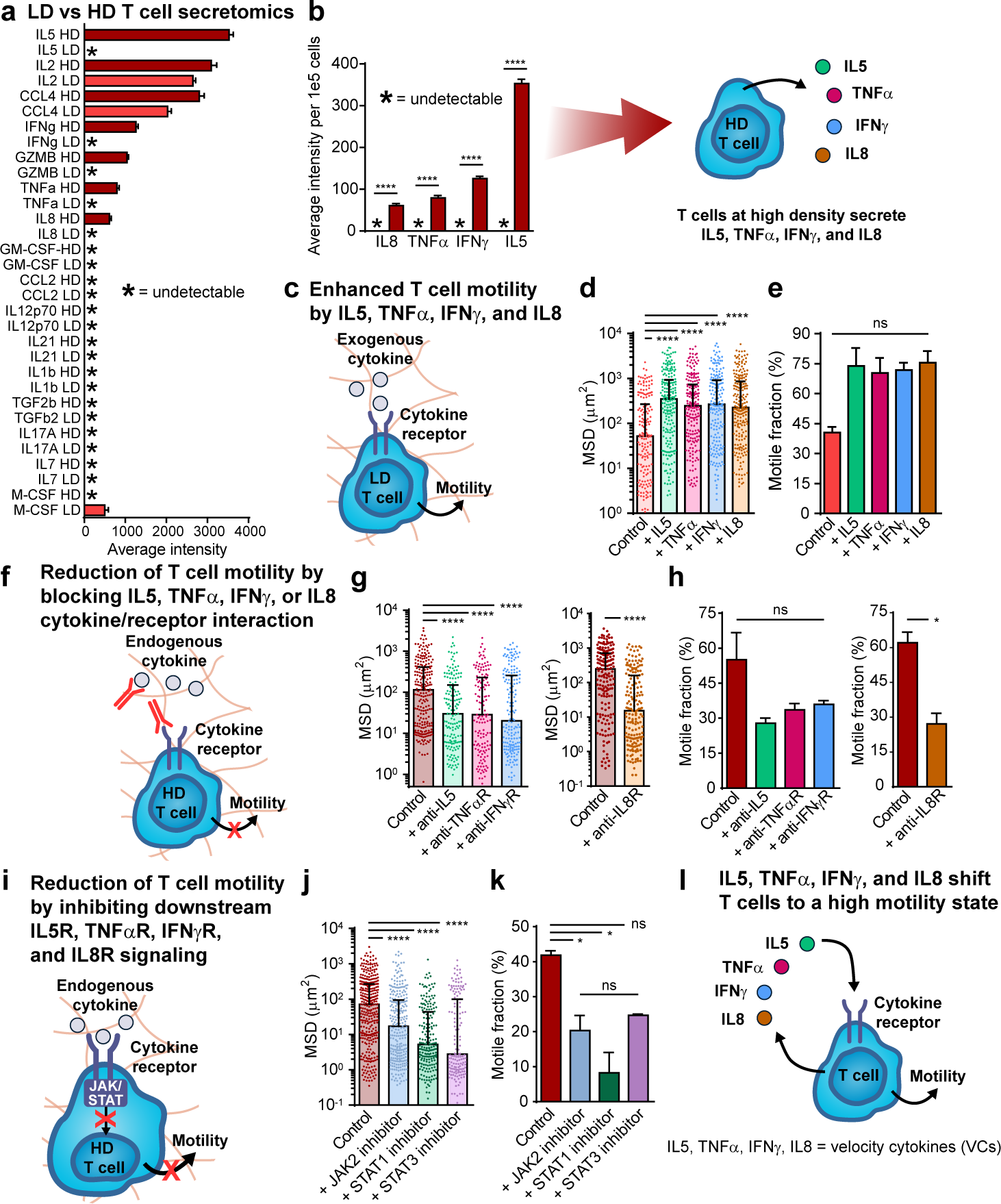
Low and high-motility states of T cell migration *in vitro* regulated by specific endogenously expressed cytokines. **a**, Molecules secreted by LD and HD T cell in 3D collagen gels incubated for 48 h. CM was extracted from the top of LD and HD T cell gels. Single-cell secretomics were run on CM to generate average intensity of a human adaptive immune cytokine panel using Isoplexis Isocode chips. **b,** Average intensity (from **a**) normalized per 10^5^ cells for the four cytokines identified (from **a**) with higher intensities for HD CM compared to LD CM. Data is plotted as mean with SEM (*n* = 2). One-way ANOVA with Dunn’s multiple comparison test was performed for statistical analysis. *\*\*\*\*P* < 0.0001. **c,** Activated LD T cells, from a new donor, in 3D collagen gels were incubated for 48 h. 100 ng/ml IL5, TNFα, IFNγ, or IL8 were then exogenously added to LD T cell gels separately and incubated for 1 h at 37 °C and 5% CO_2_. **d,e,** MSDs of T cells in the presence of IL5, TNFα, IFNg, or IL8 (**d**) and corresponding motile fractions (**e**). **f,** Activated HD T cells in 3D collagen gels were incubated for 48 h. 25 mg/ml anti-IL5, anti-TNFαR1, anti-IFNγR1, or anti-IL8R were then exogenously added separately and incubated for 1 h at 37 °C and 5% CO_2_. **g,h,** MSDs of HD T cells incubated with different functional antibodies (**g**) and corresponding motile fractions (**h**). **i,** Activated HD T cells, from a new donor, in 3D collagen gels incubated for 48 h. 2.5 mM anti-JAK2 (AG490), 60 mM anti-STAT1 (fludarabine), or 50 mM anti-STAT3 (S3I) were then exogenously added separately and incubated for 1 h at 37 °C and 5% CO_2_. **j,k,** MSDs of HD T cell gels incubated with different inhibitors (**j**) and corresponding motile fractions (**k**). **l,** T cells self-propel using velocity cytokines (VCs) IL5, TNFα, IFNγ, and IL8. For all experiments MSD, median with the SEM are plotted (*n*=2 technical replicates per biological replicate, *N*=2 biological replicates). Individual dots represent individual cells; an average of at least 80 cells per technical replicate, corresponding to up to 242 cells per technical replicate were tracked (see source data). One-way ANOVA with Dunn’s multiple comparison test was used for statistical analysis (ns = not significant, *\*P* < 0.01, *\*\*\*\*P* < 0.0001). For all experiments measuring the motile fraction, mean with SEM is plotted (*n*=2 technical replicates per biological replicate, *N*=2 biological replicates).

To understand if and how each of these cytokines modulated T cell motility, we exogenously added each recombinant cytokine separately to LD T cells in 3D collagen gels (Fig. 2c). All cytokines increased T cell motility (Fig. 2d and Extended Data Fig. 1i,j) by increasing the fraction of cells in the high-motility state (Fig. 2e and Extended Data Fig. 1k,l). Thus, all four cytokines – IL5, TNFα, IFNγ, and IL8 – can act on their own to increase T cell migration. As a control, we next inhibited the receptor/cytokine interactions of cytokines endogenously produced at detectable levels in HD T cells: IL5, TNFα, IFNγ, and IL8 using antibodies (Fig. 2f). Blocking each receptor/cytokine interaction hampered T cell migration (Fig. 2g and Extended Data Fig. 1m-p) and, correspondingly, shifted a larger fraction of T cells to a low-motility state (Fig. 2h and Extended Data Fig. 1q-t). As another control and since each of the cytokine receptors signal through JAK/STAT, we next inhibited JAK/STAT signaling in HD cells using small-molecule inhibitors (Fig. 2i), for which we observed a marked decrease in T cell motility (Fig. 2j and Extended Data Fig. 1u,v) and a shift into the low-motility state (Fig. 2k and Extended Data Fig. 1w,x).

In sum, T cells at high density produce IL5, TNFα, IFNγ, and IL8 to shift their population to a self-propelled high-motility state and subsequently increase their population-averaged migration. We termed these cytokines, velocity cytokines (VCs) (Fig. 2l).

### Velocity receptors push CAR T cells to a self-propelled high-motility state

Current cellular therapies such as CAR T suffer from inadequate infiltration into solid tumors^3^. We thus developed a migration strategy to enhance the infiltration of CAR T cells into solid tumors. Mesothelin CARs have the potential to ameliorate outcomes in many different solid tumor cancers, such as ovarian, pancreatic, and lung among others^5,6,31^. Engineering increased infiltration of mesothelin-targeted CARs has been demonstrated to enhance therapeutic outcomes in immune excluded solid tumors^16^. On this basis, we decided to enhance the migration potential of mesothelin-targeted M5CARs (M5CAR, NCT03054298 and NCT03323944) by leveraging VCs.

For our enhanced migration strategy, we ruled out overexpressing VCs in CAR T cells as all three VCs – TNFα, IFNγ, and IL8 – have been implicated in cytokine release syndrome (CRS), one of the most severe CAR T cell therapy-related toxicity^32^. We thus decided to make M5CAR T cells that were more sensitive to VCs expressed by T cells themselves or already present in their tumor microenvironment, through the overexpression of velocity receptors (VRs). These receptors bind VCs and signal through VC cognate receptor signaling domains (Fig. 3a). CAR T cells co-expressing VRs would thus exploit VCs that they secrete themselves and become self-propelled highly motile cells. We call these T cells, VR-CAR T cells.

**Fig. 3.**
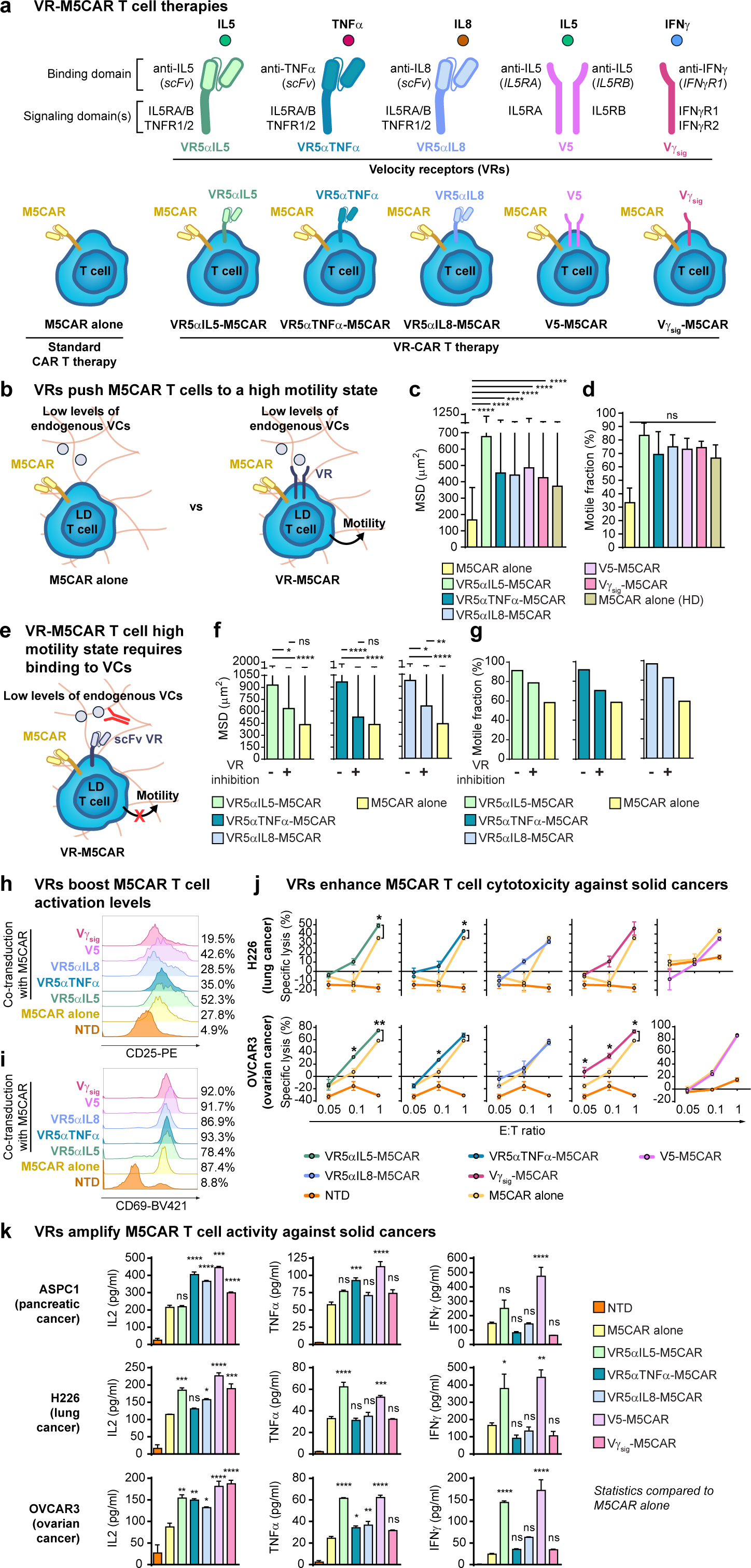
Velocity receptors turn M5CAR T cells into self-propelled highly motile cells and improve their effector functions. **a,** Design of velocity receptors (VRs) that bind VCs and signal through VC receptor signaling domains, and development of VR-M5CAR therapies. **b,** M5CAR or VR-M5CAR T cells, from a new donor, at low density (LD) were encapsulated in 3D collagen gels supplemented with 100 IU/ml IL2 and incubated for 48 h at 37 °C and 5% CO_2_ and their migration patterns were analyzed as in Fig. 1. **c**, Control HD M5CAR T cells were also encapsulated in collagen gels and incubated for 48 h and compared to LD VR-M5CAR T cells The spontaneous motion LD M5CAR, LD VR-M5CAR, and HD M5CAR T cells was monitored using time-lapsed phase-contrast microcopy for 1 h at 2-min intervals at 37 °C and 5% CO_2_. Cells were manually tracked to generate trajectories for each cell (see Extended Data Fig. 3). A custom software was used to convert trajectories into MSD for each cell at a time lag of 10 min. Each dot represents an individual cell. **d,** Fractions of T cells moving more than their own size (>R^2^ = 25 μm^2^, where R = 5 μm is the average T cell radius), as assessed from the MSDs shown in panel (**c**) (see also Extended Data Fig. 3). **e,** LD M5CAR cells expressing either VR5αIL5, VR5αTNFα, or VR5αIL8, embedded in 3D collagen gels were incubated for 48 h. 25 μg/ml anti-IL5, anti-TNFα, or anti-IL8 were then exogenously added to VR5αIL5-M5CAR, VR5αTNFα-M5CAR, or VR5αIL8-M5CAR cells, respectively, and incubated for 1 h at 37 °C and 5% CO_2_. **f,** MSDs of control LD M5CAR T cells and the different VR-M5CAR T cells shown in (**a**). T cells were imaged for 30 min at 2-min intervals (as in **c**). Cell trajectories and MSD were measured and computed (as in **c**). **g,** Motile fractions of T cells were obtained (as in **d**) for data shown in (**f**). For all experiments, MSD, median with the SEM are plotted (*n*=2 technical replicates per biological replicate, *N*=2 biological replicates for (**c**)). Individual dots represent individual cells; an average of at least 120 cells per technical replicate up to 269 cells per technical replicate were tracked (see source data). One-way ANOVA with Dunn’s multiple comparison test was used for statistical analysis (ns = not significant, *\*P* < 0.05, *\*\*P* < 0.01, *\*\*\*\*P* < 0.0001). For experiments measuring the motile fraction, mean with SEM is plotted (*n*=2 technical replicates per biological replicate, *N*=2 biological replicates) for (**d**), and mean is plotted (*n*=2 technical replicates) for (**g**). **h,** Surface expression of the activation markers CD25 and CD69, as determined by flow cytometry using PE anti-CD25 and BV421 anti-CD69. Non-transduced (NTD) T cells were included as a negative control. Surface expression of CD25 was analyzed in FlowJo on M5CAR^+^-gated T cells as determined by GFP expression. **i,** Surface expression of the activation maker CD69 was determined by flow cytometry using BV421 anti-CD69. Flow cytometric analysis was performed (as in **h**). **j,** *In vitro* T cell killing of mesothelin-expressing H226 and OVCAR3 cells at the indicated effector-to-target (E:T) ratios. Data plotted as mean ± SEM, *n* = 3. A two-tailed student’s *t* test was used for statistical analysis (*\*P* < 0.05, *\*\*P* < 0.01). **k,** Il2, TNFα, and IFNγ secretion by non-transduced (NTD), M5CAR T cells, and VR-M5CAR T cells after overnight co-incubation with mesothelin-expressing ASPC1, H226, and OVCAR3 cells. Cytokine release was measured using ELISA. Mean with SEM is shown (*n* = 3). One-way ANOVA with Tukey’s multiple comparison test was used for statistical analysis (ns = not significant, *\*P* < 0.05, *\*\*P* < 0.01, *\*\*\*P* <0.001, *\*\*\*\*P* < 0.0001).

Since there are several VCs and each VCs’ cognate receptor is a heterodimer (except for IL8 which binds to both CXCR1 and CXCR2, composed of distinct domains)^33^, we made the VR design modular. We first defined a VR as composed of four domains taken from a pool of each VCs’ cognate heterodimer receptor domains – binding, hinge, transmembrane (TM), and signaling. Next, we included in this pool of domains four VC-specific scFvs (one for each VC), and thus a CD8a hinge and CD8a TM domain to tether the scFv to the cell membrane. These were included in part to 1) allow for a single receptor in contrast to a heterodimer, and 2) yield a higher binding affinity for VCs than their respective cognate receptors. We had > 30,000 VRs we could construct (Extended Data Fig. 2a). Out of practicality, we decided to move forward with our 5 VRs to test and compare *in vitro* and preclinically *in vivo* (Fig. 3a). VR5αIL5, VR5αTNFα, and VR5αIL8 were designed to target soluble IL5, TNFα, and IL8, respectively, with corresponding anti-cyokine scFvs (refer to methods). All three were designed with CD8α hinge and transmembrane domains, and signaling domains combining both receptor chains from IL5R and TNFR (IL5RA-IL5RB-TNFR1-TNFR2) (Fig. 3a and Extended Data Fig. 2b). V5 is an overexpression of the native IL5 receptor while Vγ_sig_ has an IFNγR1 binding, hinge, and TM domain, but with signaling domains from both receptor chains (IFNγR1-IFNγR2) (Fig. 3a, and Extended Data Fig. 2b). Hence, the overall signaling domains used here were IL5R, TNFR, and IFNGR. The efficacy of IL8R signaling, as it is also a chemokine receptor, was already tested with CARs^18^.

Since all current FDA-approved CAR T cell therapies are generated by viral transduction, we generated VR-CAR T cells by co-transducing T cells with two lentiviruses: one encoding the M5CAR and the other encoding each individual VR (Extended Data Fig. 2b). VR expression was detected by antibodies using flow cytometry (Extended Data Fig. 2b and Extended Data Fig. 3a). The M5CAR was cloned into a lentiviral plasmid to express M5CAR-P2A-eGFP for flow cytometry detection (Extended Data Fig. 2b and Extended Data Fig. 3b).

As VRs were engineered to shift cells to a high-motility state, we next characterized the migration status of VR-CAR T cells to determine if these cells indeed migrated more than control CAR T cells (Fig. 3b). We found that all 5 VRs increased the migration of M5CAR T cells at LD up to levels similar or higher than M5CAR T cells at HD (Fig. 3c and Extended Data Fig. 3c,d), and shifted a higher fraction of LD M5CAR T cells to a high-motility state (Fig. 3d and Extended Data Fig. 3e,f). We determined if this observed increased migration was the result of ligand-independent tonic VR signaling in the 3 scFv-based VR-CAR cells, or if it was caused by VC binding as intended. We inhibited VC-VR binding using (separately) anti-IL5, anti-TNFα, or anti-IL8 function-blocking antibodies (Fig. 3e). With these inhibitors added, VR-M5CAR T cells exhibited hindered motility (Fig. 3f and Extended Data Fig. 3g-j) and shifted to a low-motility state (Fig. 3g and Extended Data Fig. 3k-p).

We assessed the activation status of VR-M5CAR T cells compared to M5CAR alone through the measurement of expression of standard activation markers CD25 and CD69. All VR-CAR T cells, except Vγ_sig_-M5CAR T cells, had higher CD25 expression than M5CAR T cells alone (Fig. 3h). They had roughly equal to or greater expression of CD69 than M5CAR alone (Fig. 3i). Using a standard overnight cell-killing assay, we observed that VR-M5CAR cells were cytotoxic against multiple mesothelin-expressing solid cancer cells *in vitro* at similar levels to or greater levels than M5CAR T cells (Fig. 3j). VR-CAR T cells were also either more active than or similarly active to M5CAR T cells, as determined by release of IL2, TNFα, and IFNγ in response to overnight co-incubation with multiple mesothelin-expressing solid tumor cells *in vitro* (Fig. 3k).

These results demonstrate that VRs “lock in” a self-propelled high-motility state in CAR T cells. We observed that VRs also have the added benefit of making CARs more activated, cytotoxic, and elicit stronger responses in the form of IL2, TNFα, and IFNγ production against multiple solid cancers *in vitro.* These results suggested that the performance of VR-CAR T cells in preclinical mouse models could exceed that of M5CAR T cell therapy.

### VR-CAR T cells infiltrate solid tumors in a pancreatic cancer preclinical model

The overexpression of mesothelin in about 30% of cancers^34^ makes it a promising target for the treatment of pancreatic cancer^35–37^. While mesothelin CAR T cells have been tested for this indication (NCT03323944), overall therapeutic efficacy may have been limited by poor tumor infiltration^5,6^. The vast majority of pancreatic cancers are known to be immune excluded, and therapeutic T cells are often observed around the tumor in the surrounding stroma and unable to penetrate into the tumor core^38^. However, when infiltration is observed, it is correlated with response^39^. Using the subcutaneous (s.c.) AsPC1 NSG mouse model of pancreatic cancer extensively used in previous CAR T studies^17,40,41^, we evaluated and compared the degree of infiltration and therapeutic efficacy of standard M5CARs and VR-CAR T cells. To show the utility of VR-CAR T cells, we purposefully selected a single dose (3 x 10^6^ CAR T cells) where standard M5CAR T cells alone were unable to infiltrate and control tumor growth.

As expected, control M5CAR T cells (no VRs) were detected at low levels within tumors and often surrounding the tumor, limiting their ability to reduce tumor burden (Fig. 4a and Extended Data Fig. 4a-c with additional controls). In contrast, even at this low dose, we found that several VR-M5CAR T cells infiltrated tumor cores readily and inhibited tumor growth (Fig. 4 and Extended Data Fig. 4). Not all VRs mediated a high number of M5CARs into tumors. VR5αTNFα-M5CAR and VR5αIL8-M5CAR showed similar responses to the CAR alone, with many T cells still stuck at the tumor periphery (Fig. 4b,c and Extended Data Fig. 4d,e). Interestingly, Vγ_sig_-M5CAR was detectable in the tumor core to a much higher degree (Fig. 4d and Extended Data Fig. 4f). Both VR5αIL5-M5CAR and V5-M5CAR cells infiltrated the entire tumor, edge and core, in large amounts (Fig. 4e,f and Extended Data Fig. 4g,h). Both VR5αIL5-M5CAR and V5-M5CAR inhibited tumor growth (Fig. 4e,f and Extended Data Fig. 4g,h). These qualitative assessments of CAR T cell infiltration were confirmed quantitatively using image analysis at single-cell resolution and normalized by the size of the tumor (Fig. 4g). For the best treatment, V5-M5CAR, tumor infiltration was increased by >10-fold compared to control M5CAR cells. This is an underestimate of the effect of V5 expression on CAR T infiltration, as the majority of M5CAR cells congregated at the tumor periphery.

**Fig. 4.**
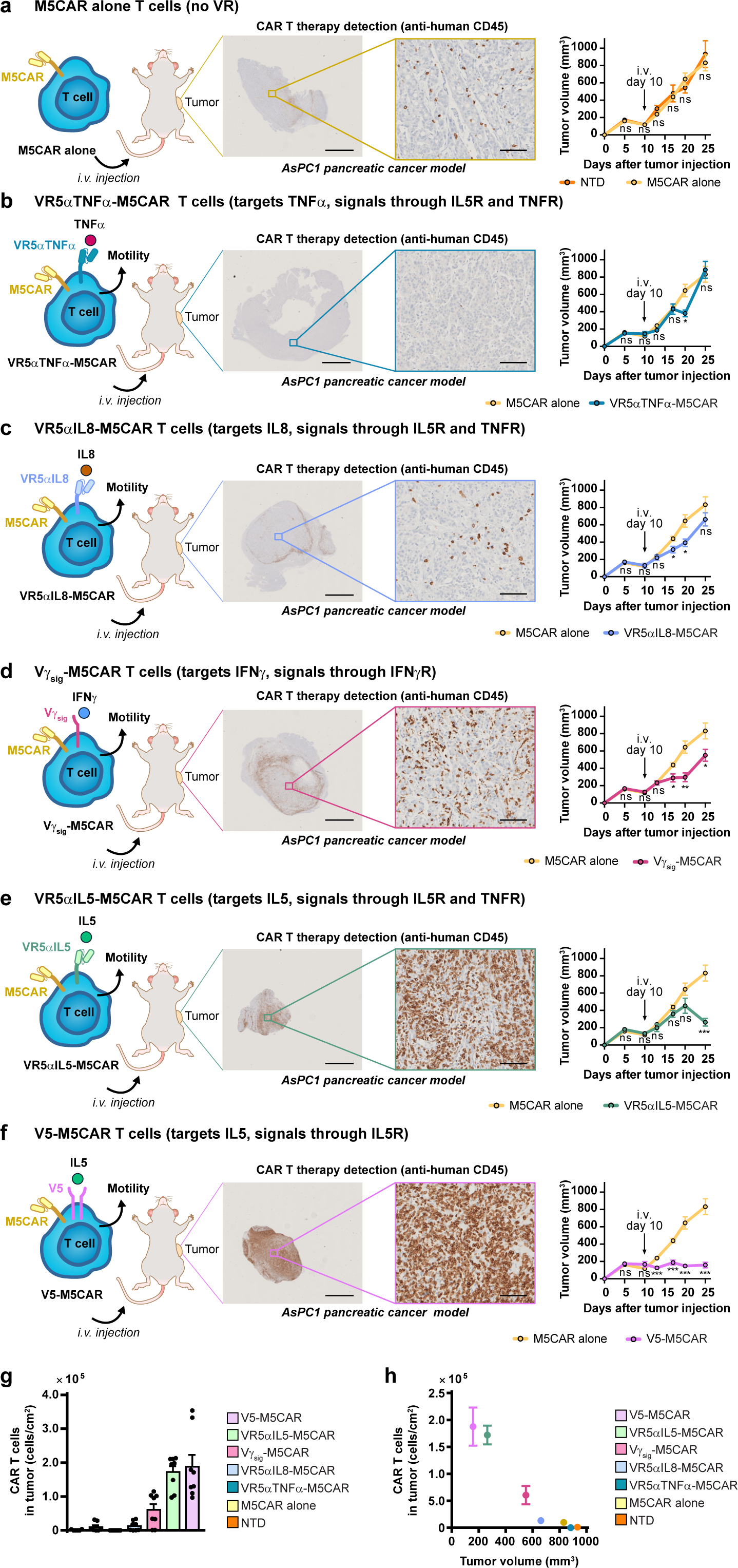
VRs greatly elevate M5CAR T cell tumor infiltration for improved controlled tumor growth in a pancreatic cancer model. **a,** 2×10^6^ ASPC1 pancreatic cancer tumors pre-mixed in 1:1 Matrigel:PBS were subcutaneously (s.c.) engrafted into 8-12 week old NSG mice. Tumor volumes were measured twice a week using digital calipers. When tumors were palpable (100 – 250 mm^3^), mice were randomized and blindly treated 10 days later with a single intravenous (i.v.) dose of 3×10^6^ NTD or M5CAR T cells. 25 days after tumor engraftment, tumors were harvested and sent for sectioning and IHC staining for human CD45 at the Johns Hopkins Oncology Tissue Services Core Facility. IHC-stained tissue slides were scanned, and images were extracted using ImageScope (middle panel). Tumor volumes were calculated as (L x W^2^) × 0.5 plotted as mm^3^ (right panel). **b,** ASPC1-bearing mice were treated with M5CAR alone or VR5αTNFα−M5CAR T cells (as in panel **a**). IHC images and tumor volumes were obtained (as in panel **a**). **c,** ASPC1-bearing mice were treated with M5CAR alone or VR5αIL8−M5CAR T cells (as in panel **a**). IHC images and tumor volumes were obtained (as in panel **a**). **d,** ASPC1-bearing mice were treated with M5CAR T cells alone or Vγ_sig_−M5CAR T cells (as in panel **a**). IHC images and tumor volumes were obtained (as in panel **a**). **e,** ASPC1-bearing mice were treated with M5CAR alone or VR5αIL5−M5CAR T cells (as in panel **a**). IHC images and tumor volumes were obtained (as in panel **a**). **f,** ASPC1-bearing mice were treated with M5CAR alone or V5−M5CAR T cells (as in panel **a**). IHC images and tumor volumes were obtained (as in panel **a**). **g,** Human CD45+ cell numbers were obtained as described in materials and methods. Tissue area was computationally obtained. Tumor infiltration of human CD45+ cells per area was calculated and plotted. **h,** Human CD45+ cells per tumor area were plotted against average tumor volume at the study endpoint. For figure panels **a-f**, IHC scale bars are 2 mm (left) and 100 μm (right). Tumor volumes are plotted as mean ± SEM (*n* = 5 mice per group, except for NTD for which *n* = 4). Two-tailed student’s *t* test was used for statistical analysis (ns = not significant, *\*P* < 0.05, *\*\*P* < 0.01, *\*\*\*P* <0.001). For calculation of cells/cm^2^ in figure panels **g,h**, *n* = 6 for NTD or *n* = 8 for VR-M5CARs, with 2 non-consecutive tissue slides stained per mouse and 3-4 mice per group; *n* = 7 for M5CAR alone, with 2 non-consecutive tissue slides stained for 3 mice and 1 tissue slide stained for 1 mouse. See Extended Data Fig. 5a for individual tumor growth curves.

We plotted the average volume of each tumor at the endpoint (Fig. 4a-f) as a function of the average number of cells that had infiltrated that tumor (Fig 4g) and found a remarkable inverse correlation (Fig. 4h): more CAR T cell infiltration mediated greater inhibition of tumor growth, which supports our original hypothesis. The inhibition of tumor growth mediated by VR-M5CAR T cells was validated using T cells from an additional donor (Extended Data Fig. 6).

We assessed potential off-tumor effects of control M5CAR T cells and VR-M5CAR cells by measuring T cell burden in non-targeted organs, including the kidney and pancreas. We also measured CAR detection in the spleen, which is a therapeutically beneficial indication of CAR expansion^42^. We found that VR-M5CARs had undetectable to minimal infiltration into the kidney and pancreas; far less than their accumulation in tumors (Extended Data Fig. 7a-g). VR5αIL5-M5CAR and V5-M5CAR cells were detected in the spleen at far higher levels than other treatments (Extended Data Fig. 7a-f). Moreover, we observed no statistically significant loss of weight in mice undergoing VR-M5CAR therapy (Extended Data Fig. 7h).

### VR-CAR T cells infiltrate solid tumors in a lung cancer preclinical model

We tested the generality of our approach by also testing VR-CAR therapies in a preclinical tumor model of lung cancer, as this is another indication where mesothelin CAR T cells could make an impact^5^. We engrafted NSG mice subcutaneously with the mesothelin-expressing H226 cell line^43–45^. For the same ineffective dose as in the pancreatic cancer model, we did not see control M5CAR cells accumulate in either the tumor or the surrounding stroma (Fig. 5a and Extended Data Fig. 8a-c with additional controls). Co-transduction with VR5αTNFα showed similar responses to M5CAR alone (Fig. 5b and Extended Data Fig. 8d). Vγ_sig_-M5CAR, VR5αIL8-M5CAR, and V5-M5CAR each formed a ring around tumor cores and led to similar tumor growth responses as M5CAR cells alone (Fig. 5c-e and Extended Data Fig. 8e-g). VR5αIL5-M5CAR therapy had the highest accumulation within tumors, which correlated with attenuated tumor growth (Fig. 5f and Extended Data Fig. 8h). CAR T cell infiltration was quantified at single-cell resolution and normalized by the size of the tumor (Fig. 5g). For the best treatment, VR5αIL5-M5CAR, tumor infiltration was increased by >300-fold compared to control M5CAR T cells.

**Fig. 5.**
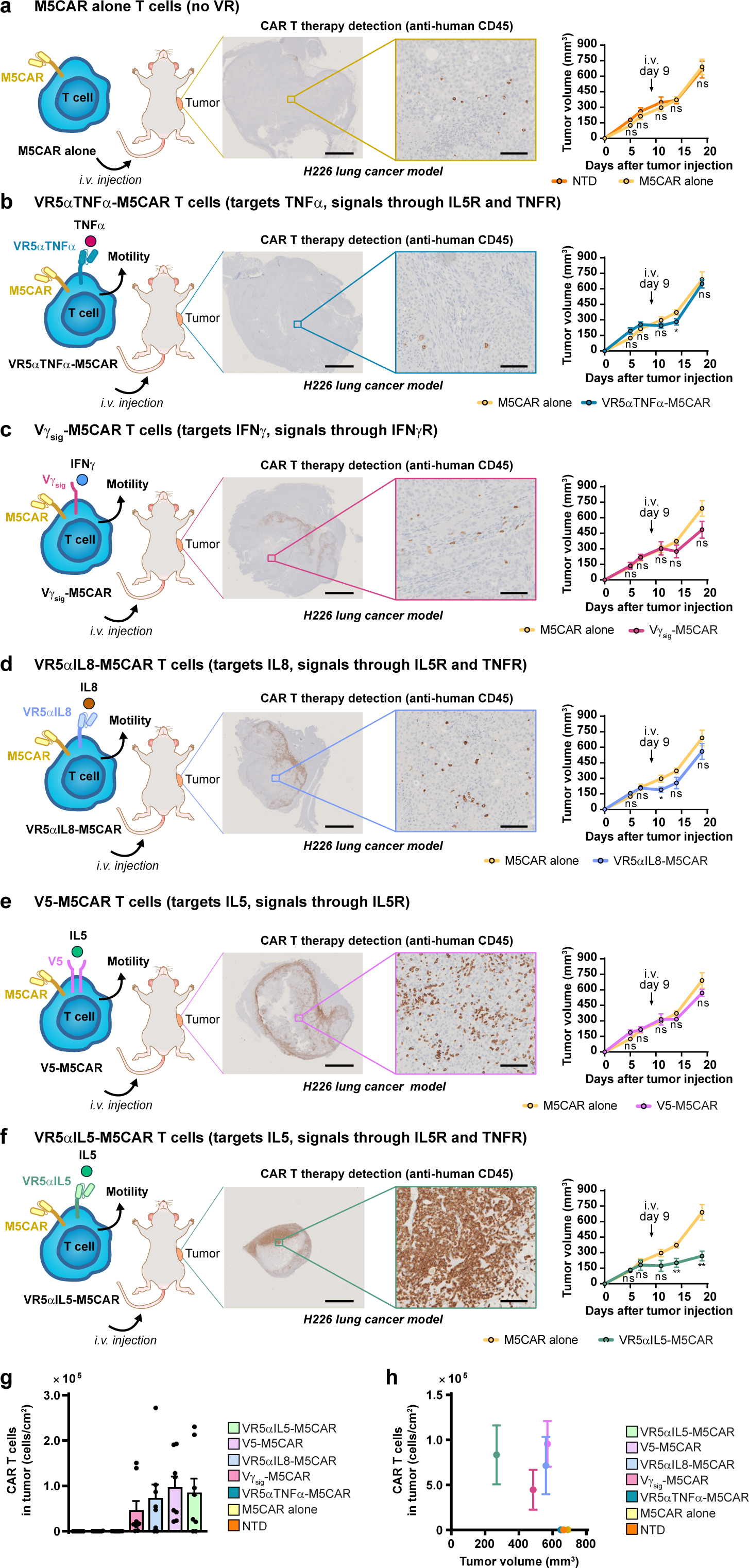
VRs improve M5CAR T cell tumor infiltration which slows tumor growth in a lung cancer model. **a,** 5×10^6^ H226 lung cancer tumors pre-mixed in 1:1 Matrigel:PBS were engrafted s.c. into 8-12 week old NSG mice. Tumor volumes were measured twice a week using digital calipers. When tumors were palpable (100 – 250 mm^3^), mice were randomized and blindly treated (9 days later) with a single i.v. dose of 3×10^6^ NTD or M5CAR T cells. 19 days after tumor engraftment, tumors were harvested and sent for sectioning and IHC staining for human CD45 at the Johns Hopkins Oncology Tissue Services Core Facility. IHC-stained tissue slides were scanned, and images were extracted using ImageScope (middle panel). Tumor volumes were calculated as (L x W^2^) × 0.5 plotted as mm^3^ (right panel). **b,** H226-bearing mice were treated with M5CAR alone or with VR5α TNFα−M5CAR T cells (as in panel **a**). IHC images and tumor volumes were obtained as in panel A. **c,** Mice bearing H226 tumors were treated with M5CAR alone or Vγ_sig_−M5CAR T cells (as in panel **a**). IHC images and tumor volumes were obtained (as in panel **a**). **d,** Mice bearing H226 tumors were treated with M5CAR alone or VR5α IL8−M5CAR T cells as in panel A. IHC images and tumor volumes were obtained (as in panel **a**). **e,** H226-bearing mice were treated with M5CAR alone or V5−M5CAR T cells as in panel A. IHC images and tumor volumes were obtained (as in panel **a**). **f,** Mice bearing H226 tumors were treated with M5CAR alone or VR5α IL5−M5CAR T cells as in panel A. IHC images and tumor volumes were obtained (as in panel **a**). **g,** Human CD45+ cell numbers were obtained as described in materials and methods. Tissue area was computationally obtained. Tumor infiltration of human CD45+ cells per area was calculated and plotted. **h,** Human CD45+ cells per tumor area were plotted against average tumor volume at the study endpoint. For figure panels **a-f**, IHC panel scale bars are 2 mm (left) and 100 μm (right). Tumor volumes are plotted as mean ± SEM (*n* = 5 mice per group, except for NTD for which *n* = 4). Two-tailed student’s *t* test was used for statistical analysis (ns = not significant, *\*P* < 0.05, *\*\*P* < 0.01). For calculation of cells/cm^2^ in figure panels **g,h**, *n* = 6 for NTD or n = 8 for M5CAR alone and VR-M5CARs, with 2 non-consecutive tissue slides stained per mouse and 3-4 mice per group. See Extended Data Fig. 5b for individual tumor growth curves.

The average volume of each tumor at the endpoint (Fig. 5a-f) was plotted as a function of the average number of cells that had infiltrated that tumor (Fig. 5g). We that more CAR T cell infiltration tended to mediate greater tumor growth inhibition (Fig. 5h), which further supports our hypothesis of an increased-migration strategy to enhance infiltration and associated therapeutic efficacy. VRs’ ability to inhibit tumor growth was validated using T cells from an additional donor (Extended Data Fig. 9a-g). Off-tumor effects was assessed for the kidney and pancreas. Accumulation of combinations with VRs was either nondetectable or miniscule (Extended Data Fig. 10a-f), especially when compared to tumor infiltration (Extended Data Fig. 10g). VR5αIL5-M5CAR and V5-M5CAR cells showed infiltration into the spleen in greater amounts than other treatments (Extended Data Fig. 10a-f). No significant loss of weight was observed in VR-M5CAR therapies compared to M5CAR alone (Extended Data Fig. 10h).

### VR-CAR T cells increase overall survival in an ovarian cancer preclinical model

Since we have established that VRs can help M5CARs accumulate within multiple solid tumors and inhibit tumor growth, we investigated if this translated into increased overall survival. To test this, we utilized an ovarian cancer preclinical mouse model (Fig. 6a), as ovarian cancers also overexpress mesothelin^5^. We subcutaneously engrafted NSG mice with the mesothelin-expressing OVCAR3 cell line^45–47^. Again, we observed that M5CAR T cells alone could not clear tumors and saw that all VR-M5CAR therapies were able to increase survival to a greater extent when compared to M5CAR alone (Extended Data Fig. 5c). Use of VR5αTNFα−M5CAR cells led to 20% overall survival after > 70 days (Fig. 6b). Co-transduction with Vγ_sig_ or V5 led to longer survival compared to M5CAR alone (Fig. 6c,d). Both VR5αIL8-M5CAR and VR5αIL5-M5CAR therapies each led to a months-long 100% overall survival (Fig. 6e,f). Compared to M5CAR alone, no significant weight loss was observed in mice treated with VR-M5CAR T cells (Extended Data Fig. 5d).

**Fig. 6.**
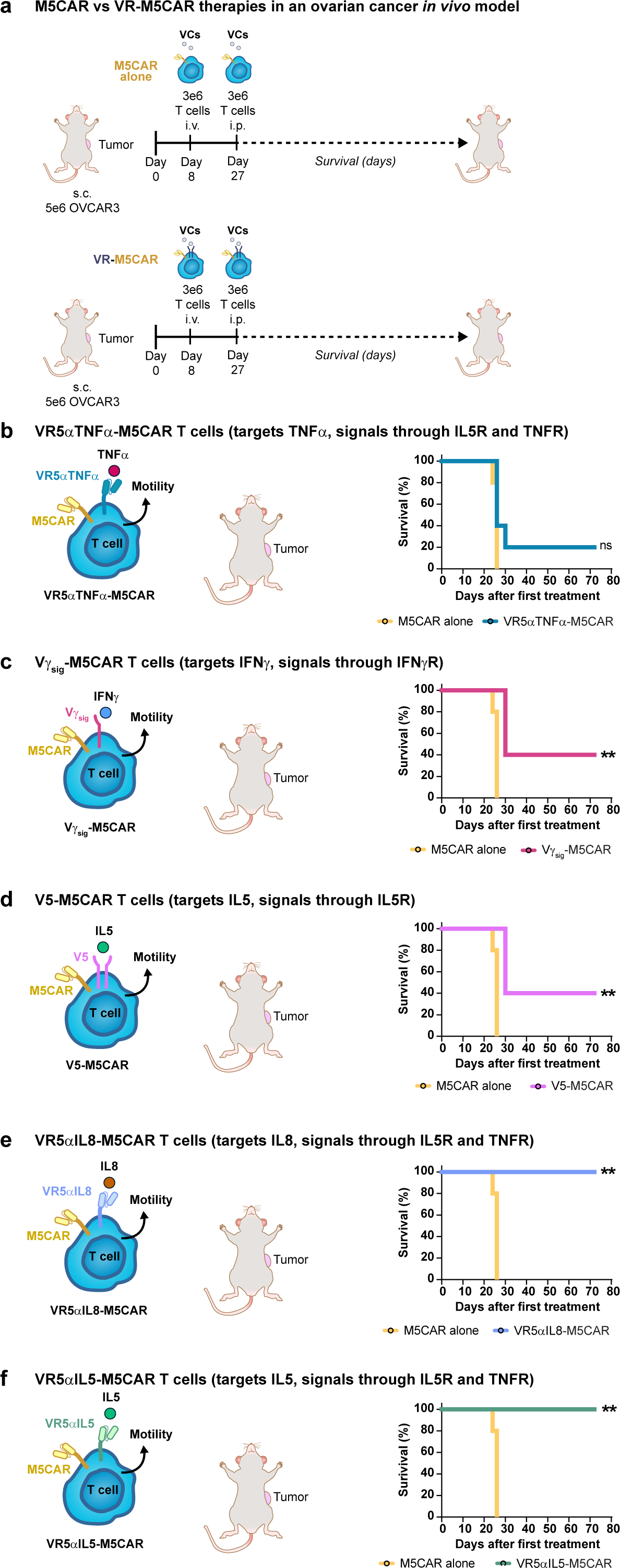
VRs-M5CAR T cells extend the overall survival of mice in an ovarian cancer model. **a,** 5×10^6^ OVCAR3 ovarian cancer tumor cells pre-mixed in 1:1 Matrigel:PBS were engrafted s.c. into 8-12 week old female NSG mice. Tumor volumes were measured twice a week using digital calipers. When tumors were palpable (100 – 250 mm^3^), mice were randomized and blindly treated (8 days later) with a single i.v. dose of 3×10^6^ M5CAR alone or VR-M5CAR T cells. Mice were treated 19 days later with 3×10^6^ M5CAR alone or VR-M5CAR T cells by intraperitoneal (i.p.) injection. Mice reached an endpoint when OVCAR3 tumors reached 750 mm^3^ in size. **b,** OVCAR3-bearing mice were treated with M5CAR alone or VR5α TNFα−M5CAR T cells. Tumor volumes were obtained as described in (**a**). Mice that reached the endpoint tumor size as described in panel (**a**) were removed from the study. Plotted are Kaplan-Meier survival curves. **c,** OVCAR3-bearing mice were treated with M5CAR alone or Vγ_sig_−M5CAR T cells. Kaplan-Meier survival curves were generated as described in panel (**b**). **d,** Mice bearing OVCAR3 tumors were treated with M5CAR alone or V5-M5CAR T cells. Kaplan-Meier survival curves were generated as described in panel (**b**). **e,** OVCAR3-bearing mice were treated with M5CAR alone or VR5α IL8-M5CAR T cells. Kaplan-Meier survival curves were generated as described in panel (**b**). **f,** Mice bearing OVCAR3 tumors were treated with M5CAR alone or VR5α IL5-M5CAR T cells Kaplan-Meier survival curves were generated as described in panel (**b**). All plots show Kaplan-Meier survival curves (*n* = 5 mice per group). Log-rank Mantel-Cox test was used for statistical analysis (ns = not significant, *\*\*P* < 0.01)

We conclude that VR-M5CAR therapies designed to increase the migration of CAR T cells in a self-propelled manner inside collagen-rich microenvironments can successfully accumulate into multiple solid tumors, control tumor growth, and increase overall survival.

## Discussion

Low levels of CAR T cell infiltration within solid tumors hinder tumor eradication in patients with solid tumors, which has greatly limited the clinical use and efficacy of CAR T therapies. This pattern is also observed with other cell therapies^48–50^, a challenge that has been typically addressed by increasing the cell therapy dose^48–50^. However, high doses lead to off-target effects and toxicity, suggesting dose-limiting toxicity. Furthermore, even when CAR Ts are used at high dose, there is still very little infiltration of CAR T cells within solid tumors^5,6^. Recent pre-clinical efforts to address poor infiltration have focused generally on making the path to the tumor easier by eradicating stromal fibroblasts surrounding tumors^17^, inducing controlled proliferation at tumor sites^16^, and trafficking CAR T cells to tumors with chemokine receptors to enhance chemotaxis^18,19^. These approaches do not directly resolve the problem of infiltration deep into solid tumor cores, which is needed to completely abrogate tumor growth.

Here we use a different strategy where we increase the migratory capacity of CAR T cells by utilizing engineering CAR T cells with receptors that take advantage of endogenously secreted cytokines that increase T-cell migration and tumor penetration. Our approach, based on our observation of two-state (high and low) motility of T cells in simple 3D collagen gels, pushes CAR T cells into a high-motility state, which facilitates CAR T cell penetration deep inside the tumor. Our analysis indicates that VR-CAR Ts can infiltrate multiple types of solid tumor in large amounts, control tumor growth, and greatly extend overall survival (Fig. 4-6). VRs that bind T cell-secreted VCs increase the fraction of migrating CAR T cells, allowing for a greater number of CAR T cells to infiltrate deep into the tumor core. This high motility observed in our VR-CAR Ts is not due to tonic signaling (Fig. 3). Our data suggests that our VR-CAR Ts are not constitutively active, as blocking the interaction of our VRs with VCs decreases the motility of our VR-CAR Ts (Fig. 3).

When we examined off-tumor infiltration, distal organ infiltration was still far lower than tumor infiltration, which is not what is seen in the clinic (Extended Data Fig. 7 and Extended Data Fig. 10). This off-tumor effect is one of the highest hurdles at the moment faced by CAR T therapeutic solutions in solid tumors^3,5,51–53^. Different methods have been designed to decrease off-tumor infiltration. A phase 1 trial of mesothelin-targeting CAR T cells electroporated with mRNA (instead of lentiviral transduction) to induce transient expression^54^, with the aim of decreasing on-target, off-tumor toxicity^5,6,54,55^. However, the transient expression appears to be too short, around 1 week^55^, to allow for enough time for CAR T cells to reach and infiltrate the tumor and generate a strong on-tumor, on-target response^5,6,54,55^. We transduced T cells lentivirally, which generated stable expression of VRs and the M5CAR. However, it would be interesting to test in future work efficacy of mRNA delivered VR-CAR constructs, as we observe high tumor infiltration in a short period following i.v. injection. Furthermore, we were able to decrease the dose of CAR T cells, and they still exhibited massive tumor infiltration despite the lower dose. It is only theorized, though likely, that M5CARs at distal sites cause antigen-targeted toxicities because it was shown that patients have low levels of mesothelin in distant organs. Howbeit, T cells with mRNA-delivered CARs in combination with VRs could exhibit substantial tumor infiltration of low-dose VR-CARs that potentially quickly eradicate tumors, while exhibiting minor distal organ infiltration that elicits a short-lived toxicity that lasts for the duration of the CAR’s transient expression. Studies could also evaluate the merit of mRNA-delivered VRs into mRNA-delivered CAR T cells; loss of VRs’ transient expression may in a sense freeze CAR T cell at high numbers in tumors and at low numbers in distant organs where after a time they would essentially revert to regular T cells. Another option for the regulation and control of VR-CAR therapies would be to introduce drug-induced suicide genes^56–58^ into VR-CAR cells to deplete them when off-tumor accumulation is observed.

It is challenging to directly compare the performance of our VR-CAR Ts to the different CAR Ts recently designed to improve infiltration, as they were tested in different mouse models and used at different doses. Nevertheless, the infiltration of VR-CAR T cells was >10 fold higher than FAP CAR T/meso CART T in a pancreatic tumor model (1.5×10^4^ cells/cm^2^ *vs*. 1.8×10^5^ cells/cm^2^)^17^. Of note, a dose of 5×10^6^ FAP CAR T cells was injected prior to 5×10^6^ mesoCAR T cells injected 15 days later, larger than the single dose of 3×10^6^ cells used here for VR-CAR T cells. The infiltration of VR-CAR T cells was also ∼180 fold higher (1×10^3^ cells/cm^2^ *vs*. 1.8×10^5^ cells/cm^2^) than CAR T cells armed with a C-X-C chemokine receptor type 6^19^, which further suggests that enhanced chemotaxis is not sufficient to enhance infiltration.

For our mouse preclinical studies, we utilized an NSG mouse model, as it is the standard model utilized to test CAR Ts that are being developed^59,60^. Utilizing this model allowed us to establish a direct comparison with previous efforts. Additionally, due to the many differences between mouse and human T cells, this model allows us to test engineered human cells, VR-CARs and CARs, that bind human ligands and human cancer cells, which are physiologically closer to human trials. The limitation of the NSG model is the lack of an immune system. A syngeneic mouse model would better evaluate the balance between tumor and off-tumor VR-CAR accumulation, as murine VR-CARs would be able to interact with murine VCs and other cells that share a similar murine background. But the resulting therapies would not be readily translatable to human subjects. In our model, no weight loss or toxicity was detected in any of our treatments. Toxicity assessments would be preferably conducted in syngeneic mouse models. Something to note is that the only cells producing VCs that our therapies would bind are the T-cell therapies themselves. We hypothesize that a higher infiltration of VR-CAR T cells would be reached in human patients as our engineered VR-CAR T cells would be able to utilize the cytokines secreted by the solid tumors and cells in the TME^61–65^.

In summary, we have engineered CAR T cells that sustain a high migratory phenotype that facilitates infiltration in solid tumors. Though we tested our VR technology in M5CAR T cells, it is of no technical limitation to co-express our VRs with any other CAR. We anticipate that the greatest utility of VRs is their combination with tumor-specific targeting receptors that only recognize neoantigens. This would eliminate solid organ on-target off-tumor toxicity and the only toxicities that arise would be those similar to what ensues from hematologic cancer targeting CAR T cell therapies, such as neurotoxicity and cytokine release syndrome which are manageable. Neoantigens can be targeted with CAR or TCR technologies. Though it is likely, what is still unknown is whether VRs are only compatible with CAR T cells or if they can also be used with TCR T cells.

## Material and Methods

### Human cancer cells

OVCAR3 (high-grade ovarian serous adenocarcinoma), AsPC1 (pancreatic ductal adenocarcinoma), H226 (squamous cell lung carcinoma) cell lines were obtained from ATCC. OVCAR3 cells were maintained in RPMI supplemented with 10% FBS, 1% penicillin/streptomycin, 200 mM L-glutamine, 1% HEPES, and 1% sodium pyruvate. AsPC1 cells were maintained in RPMI supplemented with 20% FBS, 1% penicillin/streptomycin, and 1% HEPES. We maintained H226 cells in RPMI supplemented with 10% FBS and 1% penicillin/streptomycin. For *in vitro* co-culture cytotoxicity studies, all cell lines were lentivirally transduced to stably express luciferase-P2A-mCherry. Following transduction, all cell lines underwent FACS cell sorting using a Sony SH800 cell sorter.

### Donor primary human T cells

Primary human T cells were obtained from fresh identity-unlinked leukapheresis packs (LPs) from healthy donors who underwent same-day apheresis through the Anne Arundel Medical Center (Annapolis, MD, USA), which resulted in fresh leukapheresis packs (LPs). Freshly delivered LPs were immediately processed to extract PBMCs using Ficoll density gradients. PBMCs subsequently underwent negative selection for the isolation of pan T cells, or CD4 or CD8 T cell subsets for CAR manufacturing using STEMCELL negative selection isolation kits. T cells were maintained in X-Vivo 15 media supplemented with 5% Human AB serum, 200 mM L-glutamine, and 1% penicillin/streptomycin – after which, they were referred to as complete X-Vivo. For density-dependent migration studies (Fig. 1b-g and Fig. 2a-l), T cells were cultured in serum-free Immunocult media (STEMCELL) supplemented with 100 IU/ml IL2.

### Cellular therapy manufacturing

VR therapy vectors and the CAR therapy vector were first designed through Gibson Assembly molecular cloning (HiFi DNA Assembly Cloning Kit, NEB) using the backbone pSLCAR-CD19-BBz (Addgene). VR nucleic acids were synthetically constructed as indicated (refer to Fig. 3a and Extended Data Fig. 2a-b), and were cloned into the backbone excluding the backbone’s native CD19 CAR-P2A-GFP sequence to select for positive cells downstream through the use of surface staining with FACS antibodies for VRs and subsequent detection by flow cytometry. The M5CAR nucleic acid sequence was cloned into the backbone prior to the P2A-GFP sequence to select positive cells downstream by GFP using flow cytometry. VR therapy lentiviral particles and CAR therapy lentiviral particles were obtained by transfecting 293T cells with vectors for VRs or the M5CAR. 293T cells were obtained from ATCC and maintained in either DMEM supplemented with 10% FBS or Opti-MEM supplemented with 5% FBS. Transfection was performed using GeneJuice, (Millipore Sigma) and lentiviral particles were concentrated in Lenti-X Concentrator (Lonza).

VR, CAR, and VR-CAR therapies were manufactured as follows. CD4 and CD8 T cells were isolated using STEMCELL negative selection kits and were mixed 1:1^66–70^. They were then activated overnight by CD3/CD28 Dynabeads (Invitrogen) in complete X-Vivo media supplemented with 100 IU/ml IL2 (Peprotech). For VR alone or M5CAR alone therapies, activated cells were subsequently lentivirally transduced overnight on Retronectin-coated plates, after which their positivity was checked by flow cytometry. For the VR-CAR cells, after overnight activation, T cells were transduced on Retronectin-coated plated with VR therapy lentivirus overnight and checked for VR positivity by flow as follows: V5 stained for IL5RA; Vγ_sig_ stained for IFNγR1; VR5α IL5, VR5α IL8, and VR5α TNFα stained for F(ab)_2_. VR T cells were then transduced overnight on Retronectin-coated plates with M5CAR therapy lentivirus and checked for M5CAR positivity by flow cytometric detection of GFP. VR-M5CAR, M5CAR, or non-transduced T cells were expanded for a total of 11 days^71^. Dynabeads were magnetically removed using Dynamags (Invitrogen) and frozen in liquid nitrogen or maintained in culture^51,72,73^ in complete X-Vivo media supplemented with 100 IU/ml IL2.

### 3D migration

T cells in complete X-Vivo media supplemented with 100 IU/ml IL2 were suspended in a 2 mg/ml type I collagen (Corning) solution. For density-dependent migration studies, T cells in Immunocult media were activated with Immunocult CD2/CD3/CD28 expansion solution (STEMCELL) supplemented with 100 IU/ml IL2. T cells were then suspended in 2 mg/ml type I collagen (Corning) and a 0.22 μm-filtered reconstitution buffer that consisted of HEPES, HBSS, and sodium bicarbonate in milliQ-purified water. The T cell-collagen solution was solidified in a 37 °C and 5% CO_2_ incubator for 1 hr. Immunocult media supplemented with 100 IU/ml IL2 was then added on top of the T cell-collagen gels. The gels were then incubated for 48 hrs and imaged for 1 hr on a Nikon Ti2 microscope equipped with a live-cell system maintained at 37 °C and 5% CO_2_. Time-lapse images were constructed into time-lapse movies using Nikon Elements, and cell positions were tracked frame-by-frame to generate cell trajectories using Metamorph. Analysis of cell trajectories to pull out multiple parameters to characterize cellular motility was performed in Matlab (MathWorks) following the principles and equations previously reported^13,74^.

### T cell secretomics

LD and HD T cells were cultured in 3D collagen gels, as described above, for 48 hrs in a 37 °C and 5% CO2 incubator. Conditioned medium was collected from the top of 3D T cell-collagen gels and filtered through a 0.45 μm filter. Filtered conditioned medium was added to Isoplexis microchips.

### 2D *in vitro* cytotoxicity

For T cell cytotoxicity, luciferase-expressing solid tumor cell lines were incubated at 37 °C and 5% CO_2_ overnight. Subsequently, T cells at indicated E:T ratios were added on top of the cancer cells and incubated overnight. BrightGlo solution (Promega) was then added to the wells and luminescence was read on a SpectraMax plate reader (Molecular Devices) to obtain relative light units (RLUs). Specific lysis was calculated using the RLUs as 100% * (Alive – Condition)/(Alive – Dead).

### Flow cytometry immunophenotyping

For immunophenotyping of VR-transduced T cells, 50 uL overnight-transduced T cells were collected, washed three times at 4 °C in cold FACS buffer (PBS supplemented with 5% FBS and 0.1% sodium azide), and surfaced stained with an anti-F(ab)_2_ (VR5αIL5, VR5αTNFα, and VR5αIL8), anti-IL5RA (V5), or anti-IFNγR1 (Vγ_sig_) antibody in FACS buffer at 4 °C for 30 min. Cells were then washed three times at 4 °C in FACS buffer before analysis by flow cytometry. Data was collected using Sony SH800 and was analyzed using FlowJo (BD). For M5CAR detection, T cells were collected and washed three times in FACS buffer and analyzed for GFP expression using flow cytometry.

For activation marker expression immunophenotyping, T cells were collected, washed three times at 4 °C in cold FACS buffer, and surface stained with an anti-CD25 or anti-CD69 antibody at 4 °C for 30 min. Cells were washed three times at 4 °C in cold FACS buffer. Surface-stained T cells were then analyzed by flow cytometry using Sony SH800. All data were analyzed using FlowJo software.

### Cytokine assays

To measure T cell activity, AsPC1, OVCAR3, or H226 cells were incubated in overnight at 3e3 cells/well in 96 well plates. The following day, NTD, M5CAR, or VR-M5CAR T cells in were added to cancer cells at 1:1 E:T ratios and incubated overnight. 100 μL of supernatant was then carefully removed from each wells using a multichannel pipette. Supernatants were analyzed for IL2, TNFα, or IFNγ by ELISA (Invitrogen) according to the manufacturer’s protocol.

### Pre-clinical mouse models

All animal experiments were approved by the Institutional Animal Care and Use Committee (IACUC) of the Johns Hopkins University (Protocol MOE23E29) and were performed in an Association for the Assessment and Accreditation of Laboratory Animal Care (AALAC) accredited facility. The housing group was five at maximum for all mice groups. Mouse studies were performed in 8–12-week-old NOD-*scid* IL2Rgamma^null^ (NSG) female and male mice (Jackson Laboratory). AsPC1 (2×10^6^) or H226 (5×10^6^) human cells were subcutaneously injected into the right flank of each mouse with an equal volume of Matrigel (BD) to PBS. After tumors became palpable at roughly 100 – 250 mm^3^, mice were injected by tail vein (once) with either 100 μl PBS or non-transduced T cells, M5CAR T cells, or the indicated VR-M5CAR T cell therapies in 100 μl PBS. Tumors were measured using digital calipers approximately twice per week and mice were weighed at the end of each experiment. Tumor volumes were calculated using the following formula: volume: (*L x W*^2^) *× 0.5*. Tumors and organs were harvested at the indicated days. Half of the tissues were embedded in paraffin for IHC staining and the other half was flash-frozen.

### Survival Mouse Studies

The OVCAR3 (5×10^6^) cancer cell line was subcutaneously injected into the right flank of female NSG mice. After tumors became palpable mice were injected by tail vein with the respective treatment. Mice received a second dose after 19 days intraperitoneally. Tumors were measured twice a week as described above. Mice were sacrificed when they reached 750 mm^3^, and tumors and organs were collected.

### Immunohistochemistry (IHC)

Mouse tumors and organs were fixed in formalin and subsequently paraffin-embedded and sectioned. Paraffin-embedded sections were blindly stained for human CD45 to measure human T cell infiltration into tumors and organs by The Johns Hopkins Oncology Tissue Core. Histological slides were scanned using Hamamatsu NanoZoomer S210 scanner at a 20x magnification (0.5 μm/pixel). Quantification was done by a computational biologist who was blinded to the data.

### Immunohistochemistry quantifications

Scanned histological images were downsampled to 10x magnification (1 μm/pixel) and color deconvolution was used to extract the IHC CD45 positive locations to an independent image channel. Intensity peaks were detected on the isolated CD45+ channel and spatial coordinates were collected (x, and y coordinates). Assessment of tumor CD45+ cell infiltration was performed by manual masking of each individual tumor location.

### Statistical Analysis

All statistical tests were performed using GraphPad Prism. All migration MSD analyses used either the nonparametric Mann-Whitney *U*-test for two groups, or for multiple comparisons the Kruskal-Wallis ANOVA with Dunn’s post-test. All other analyses were performed using either the student’s *t*-test for two groups, or the ordinary one-way ANOVA with Tukey’s post-test for comparison of multiple groups. The Log rank Mantel-Cox test was used for Kaplan-Meier survival curves.

## Acknowledgements

The authors acknowledge the following sources of support: U54AR081774; U54CA268083; UG3CA275681; T32CA153952 from the National Cancer Institute. The authors thank colleagues Bert Vogelstein, Ken Kinzler, Nick Papadopoulos, Max Konig and their students and fellows for fruitful feedback on this work.

