## Extended data figures for "Engineering self-propelled tumor-infiltrating CAR T cells using synthetic velocity receptors"

**Extended Data Fig. 1**

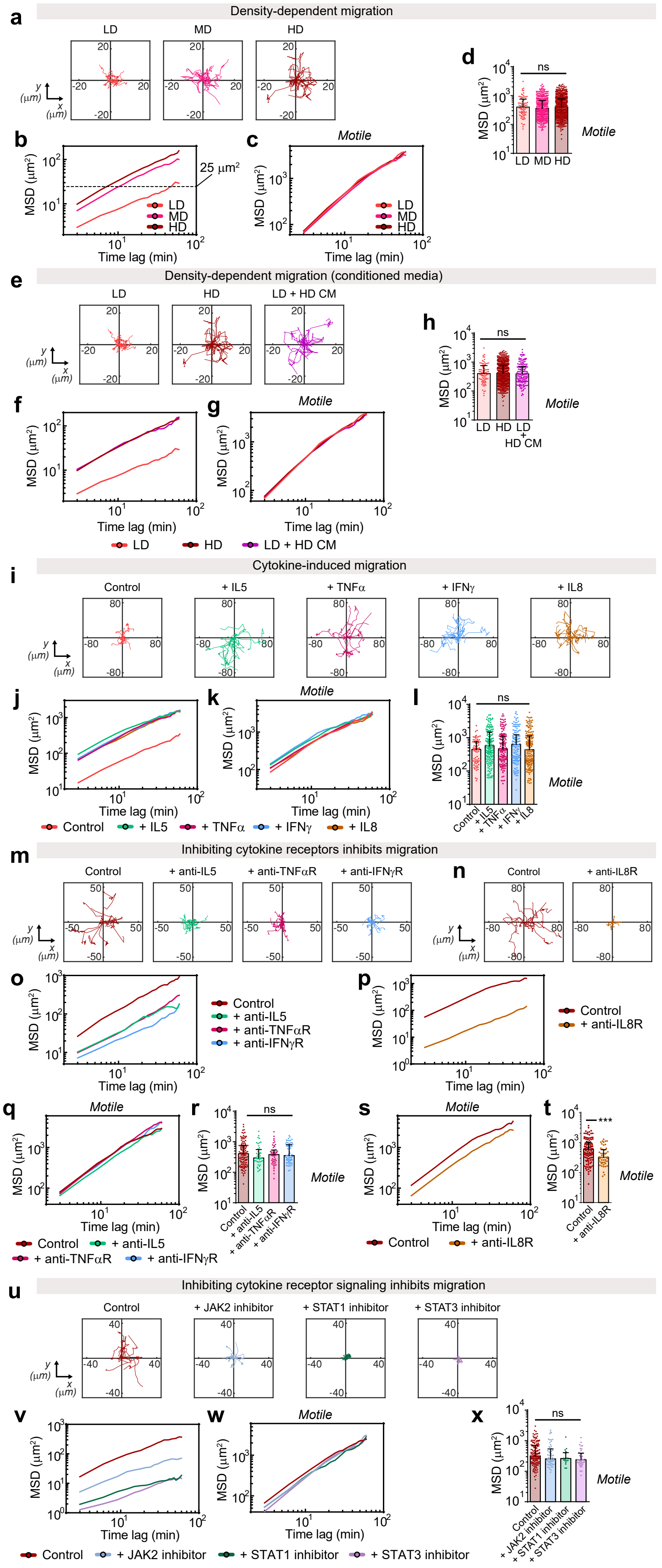

**Extended Data Fig. 1 | a,** LD, MD, and HD T cells were encapsulated in 3D collagen gels and their motility was monitored as described in Fig. 1b-d. A custom Matlab software was used to randomly select 10 trajectories per condition. Plotted are x-y trajectories (μm). **b,** Median T cell MSD was calculated and plotted versus time lag from all cell trajectories. **c,** T cells that moved more than their own size (>$R^2$ = 25 μ$m^2$, where R ~ 5 μm is the average T cell radius) in an hour of tracking were extracted and the median MSD was calculated and plotted versus time lag (min). **d,** A Matlab script was coded to extract the percentage of motile T cells MSDs at a time lag of 9 min from the MSDs in panel (**c**). **e,** LD, HD, and LD + HD T cells were encapsulated in 3D collagen gels and imaged as described in Fig. 1e-g. Ten randomly selected trajectories displayed per condition. **f,** Median T cell MSDs for the indicated conditions. **g,** MSDs of motile T cells were extracted as described in panel (**c**). **h,** MSDs of motile T cells at a time lag of 9 min from data in panel (**g**). **i,** LD T cells in 3D collagen gels were treated with exogenous cytokines and their motility was monitored as described in Fig. 2c-e. Ten randomly selected trajectories (μm) per condition are shown. **j,** Median T cell MSD was calculated and plotted for the indicated conditions. **k,** MSDs of motile T cells were extracted as described in panel (**c**). **l,** MSDs of motile T cells at a time lag of 9 min, computed from MSDs, as described in panel (**c**). **m,** HD T cells in 3D collagen gels were treated with indicated antibodies and their motility was monitored as described in Fig. 2f-h. Ten randomly selected x-y trajectories (μm) per condition are shown. **n,** HD T cells in 3D collagen gels were treated with an anti-IL8R antibody and their motility was monitored as described in Fig. 2f-h. Ten randomly selected x-y trajectories (μm) per condition are shown. **o,** Median T cell MSDs for the indicated conditions. **p,** Median T cell MSDs for HD control T cells or HD T cells treating with an anti-IL8R antibody. **q,** Motile MSDs were extracted and plotted in Matlab as described in panel (**c**). **r,** Matlab was used to extract and plot motile T cell MSDs at a time lag of 9 min from the MSDs in panel (**q**). **s,** MSDs of motile T cells were extracted as described in panel (**c**). **t,** MSDs of T cells at a time lag of 9 min shown in panel (**s**). **u,** HD T cells in 3D collagen gels were treated with indicated JAK/STAT inhibitors and their motility was monitored as described in Fig. 2i-k. Ten randomly selected trajectories per condition are shown. **v,** Median MSDs of T cells for the indicated conditions. **w,** MSDs of motile T cells were extracted as described in panel (**c**). **x,** MSDs of T cells at a time lag of 9 min computed from the MSDs in panel (**w**).

All plots showing MSD *vs*. time lag are plotted using the median MSD. For all experiments measuring MSD at a time lag of 9 min, median with the SEM is plotted ($n$=2 technical replicates per biological replicate, $N$=2 biological replicates). Individual dots represent individual cells; an average of at least 80 cells per technical replicate, up to 989 cells per technical replicate were tracked (see source data). One-way ANOVA with Dunn's multiple comparison test was used for statistical analysis (ns = not significant, ***$P < 0.001$).

**Extended Data Fig. 2**

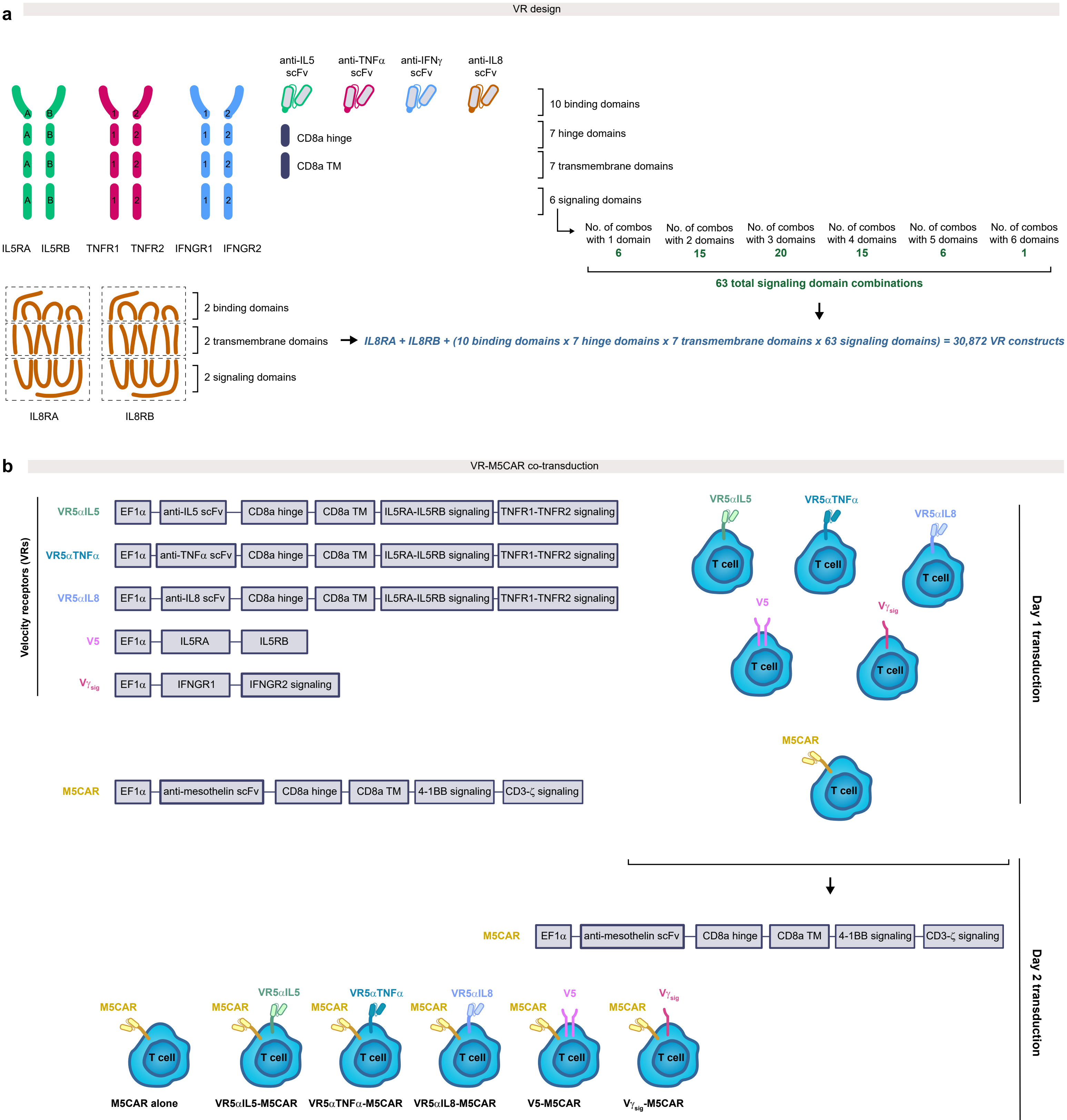

**Extended Data Fig. 2 | a,** VR design. Calculated are the number of VR constructs obtained from the indicated combination of domains. **b,** VR-M5CAR T cell co-transduction scheme.

**Extended Data Fig. 3**

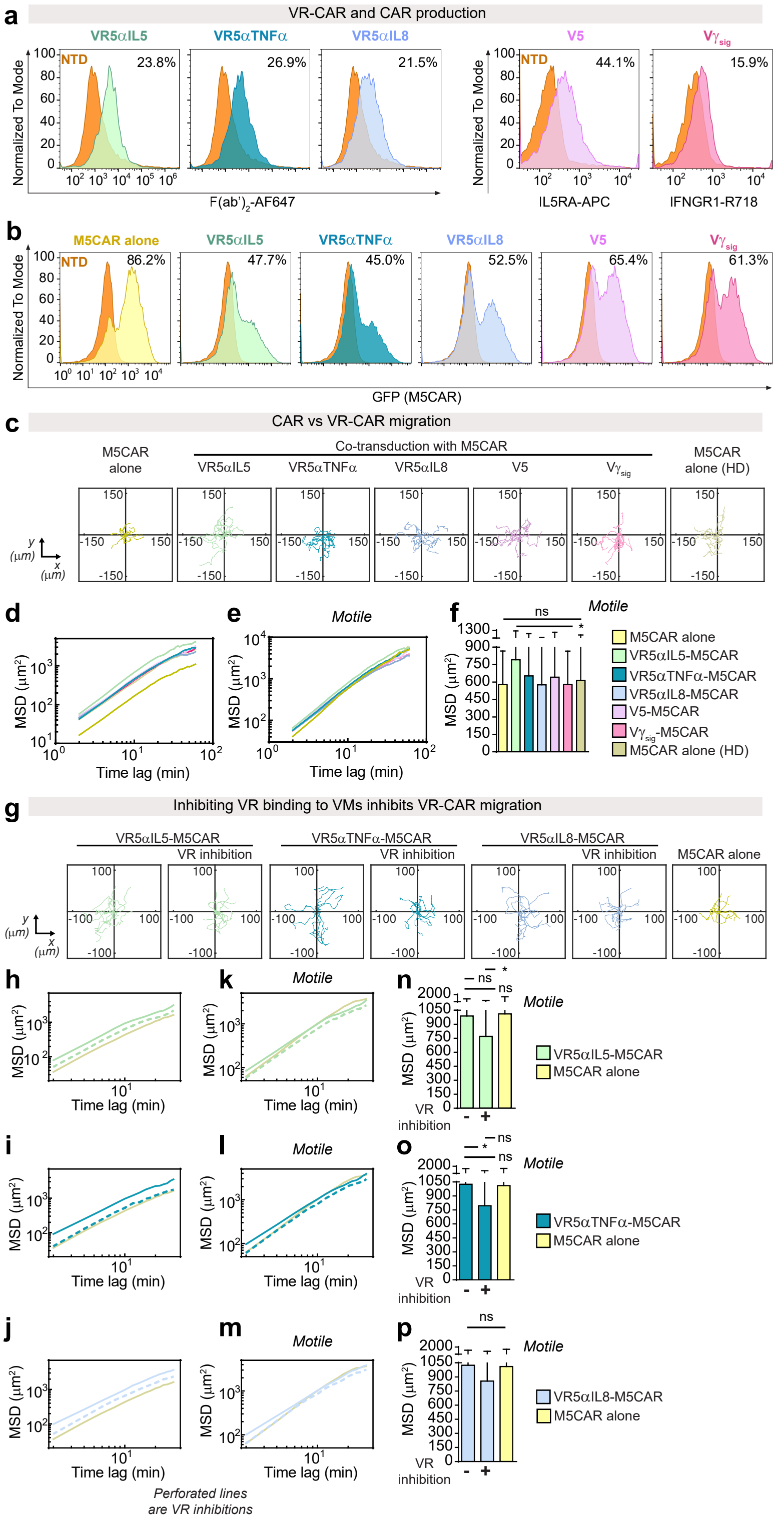

**Extended Data Fig. 3 | a,** CD4+ and CD8+ primary human T cells were isolated from Ficoll-extracted PBMCs from human donor leukopaks and mixed 1:1. T cells were then activated overnight with CD3/CD28 Dynabeads and 100 IU/ml IL2. The following day, lentiviruses bearing plasmids with transgenes that encode for the specified VRs (see Fig. 3a) were incubated on Retronectin-coated plates for 4 hrs at 37 °C and 5% $CO_2$. Activated T cells were then added to lentivirus-bound Retronectin coated wells and incubated overnight. VR surface expression was detected the next day by flow cytometry after staining with anti-F(ab)$_2$ antibodies for the VRs indicated, anti-IL5RA for V5, or anti-IFNGR1 for $V\gamma_{sig}$. Non-transduced (NTD) T cells were used as a negative control. FlowJo was used to analyze the data and plot histograms normalized to mode. **b,** The day following VR transduction, T cells were transduced overnight with M5CAR-P2A-eGFP (see Fig. 3a) on lentivirus-bound Retronectin-coated plates. The next day, M5CAR expression was measured by GFP detection using flow cytometry. Non-transduced (NTD) T cells were used as a negative control. Data was analyzed and histograms normalized to mode were plotted in FlowJo. **c,** LD M5CAR, LD VR-M5CAR, and HD M5CAR T cells were encapsulated in 3D collagen gels and their motility was monitored as described in Fig. 3b-d. Ten randomly selected trajectories per condition were selected. **d,** Median MSD of T cells vs. time lag. **e,** T cells that moved more than their own size ($>R^2 = 25\ \mu m^2$, where $R = 5\ \mu m$ is the average T cell radius) in an hour of tracking were extracted in Matlab and median MSD was calculated and plotted versus time lag. **f,** MSDs of motile T cells at a time lag of 10 min, as described in panel (**e**). **g,** LD M5CAR or LD VR5αIL5-M5CAR, VR5αTNFα-M5CAR, or VR5αIL8-M5CAR cells were suspended in 3D collagen gels, treated with antibody inhibitors, incubated, and their motility was monitored as described in Fig. 3e-g. Ten randomly selected trajectories (mm) per condition are shown. **h-j,** Median T cell MSD was calculated and plotted using a custom Matlab software. **k-m,** Motile MSDs were extracted and plotted in Matlab (as described in panels **h-j**). **n-p,** A custom Matlab script was used to extract and plot motile T cell MSDs at a time lag of 10 min from the MSDs in panels (**k-m**).

All plots showing MSD vs time lag are plotted using the median MSD. For all experiments measuring MSD at a time lag of 10 min, median with the SEM is plotted ($n$=2 technical replicates per biological replicate, $N$=2 biological replicates for (**f**) and $N$=1 biological replicate for panel **n-p**). Individual dots represent individual cells; an average of at least 120 cells per technical replicate up to 269 cells per technical replicate were tracked (see source data). One-way ANOVA with Dunn's multiple comparison test was used for statistical analysis (ns = not significant, $*P < 0.05$).

**Extended Data Fig. 4**

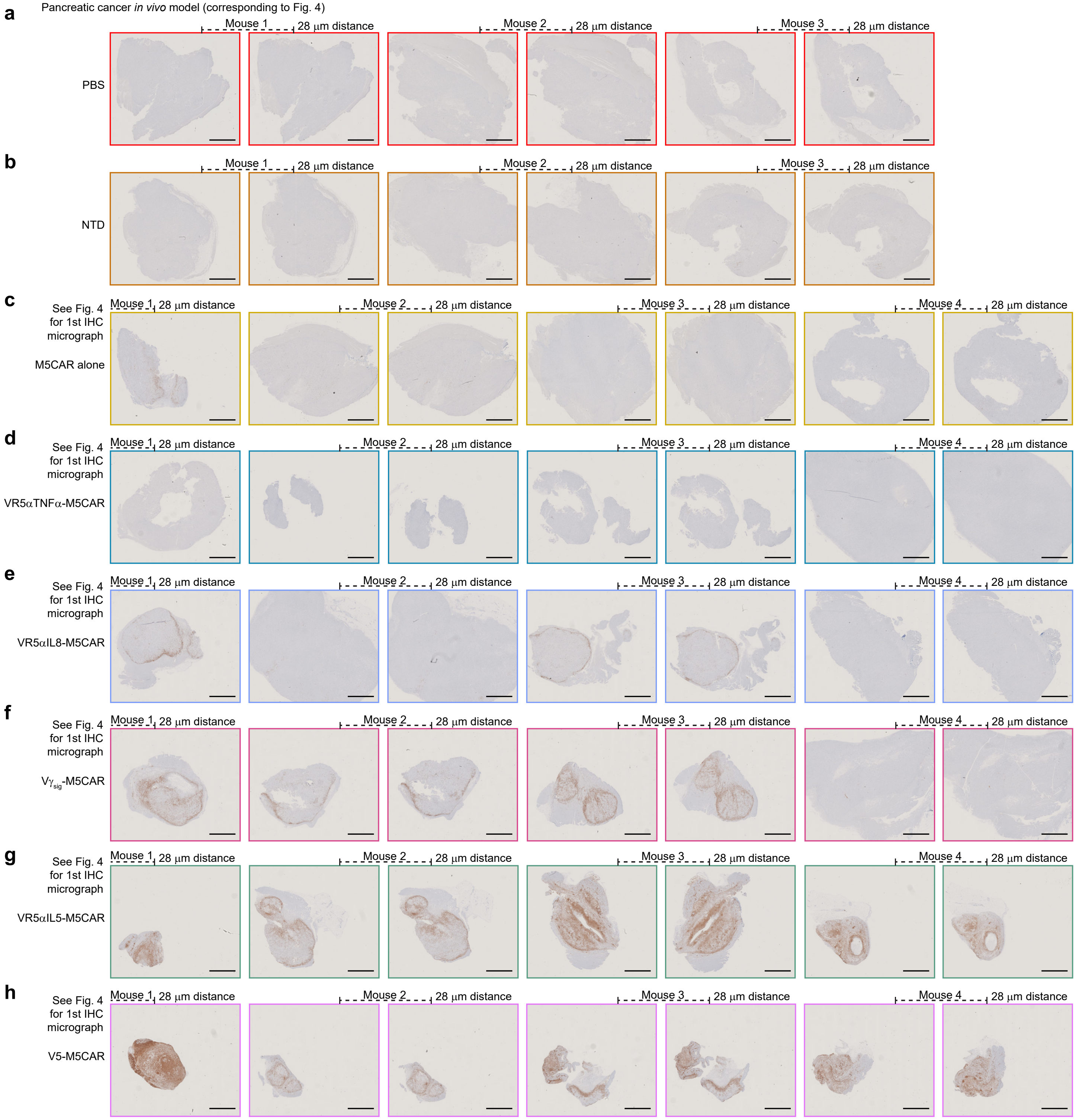

**Extended Data Fig. 4 | a-h**, CD45+ IHC micrographs of tumors from ASPC1-bearing mice (corresponding to Fig. 4) treated with PBS (**a**), NTD T cells (**b**), M5CAR alone T cells (**c**), VR5αTNFα-M5CAR T cells (**d**), VR5αIL8-M5CAR T cells (**e**), V$\gamma_{sig}$-M5CAR T cells (**f**), VR5αIL5-M5CAR T cells (**g**), or V5-M5CAR T cells (**h**). Scale bars in IHC micrographs are 2 mm.

**Extended Data Fig. 5**

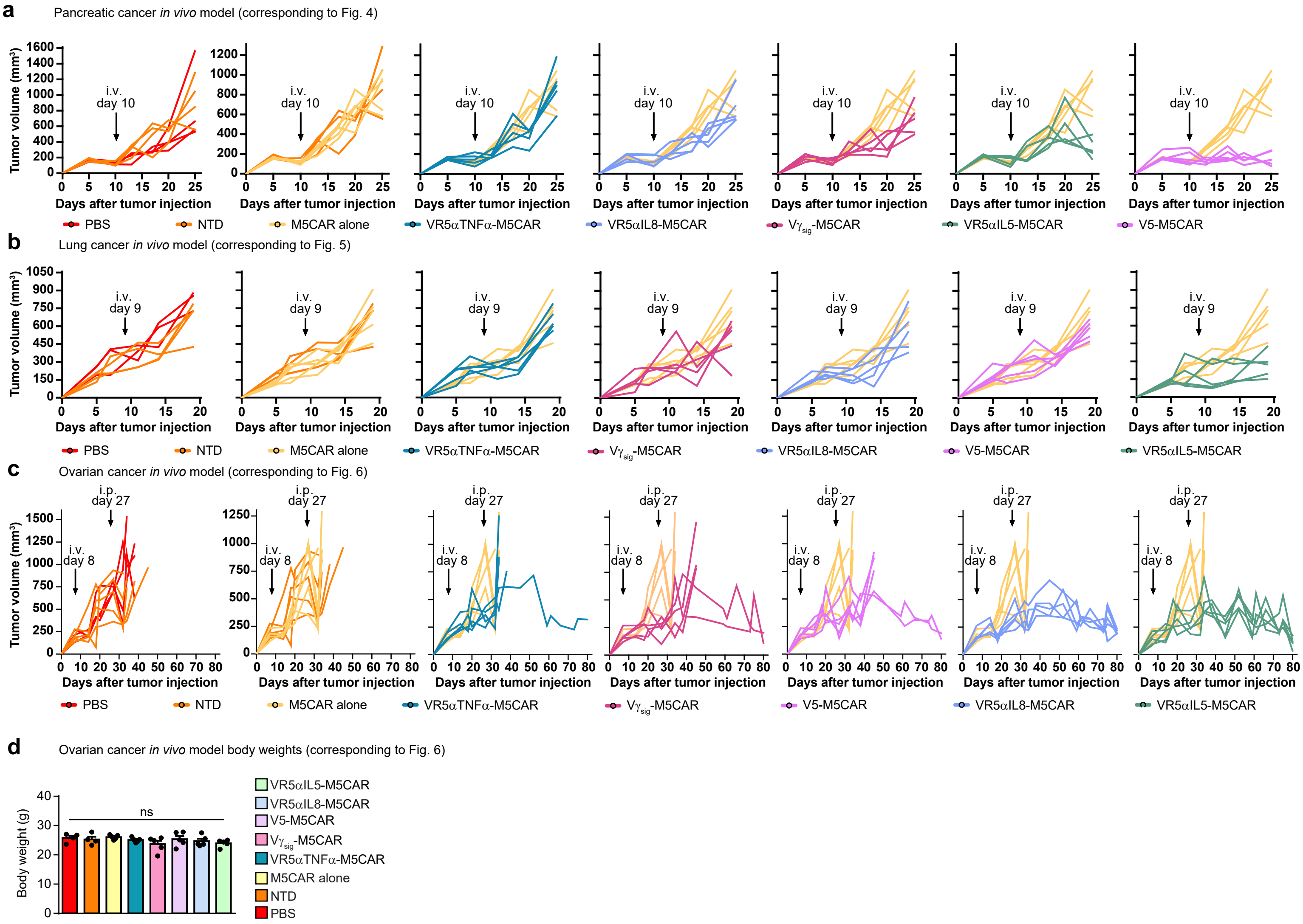

**Extended Data Fig. 5 | a-c**, Tumor growth of individual mice corresponding to ASPC1-bearing mice (as in Fig. 4) (**a**), H226-bearing mice (as in Fig. 5) (**b**), and OVCAR3-bearing mice (as in Fig. 6) (**c**) with indicated treatments. **d**, Mice were weighed at the end of the study. For **a-c**, $n = 5$ for all treatment groups, except for PBS and NTD for which $n = 4$ (**a**). $n = 5$ for all treatment groups, except for NTD for which $n = 4$ and PBS for which $n = 3$ (**b**). $n = 5$ for all treatment groups, except for PBS for which $n = 4$ (**c**). For **d**, body weights are plotted as mean ± SEM ($n = 5$ mice per group, except for PBS and NTD for which $n = 4$). Ordinary one-way ANOVA with Tukey's multiple comparison test was used for statistical analysis (ns = not significant).

**Extended Data Fig. 6**

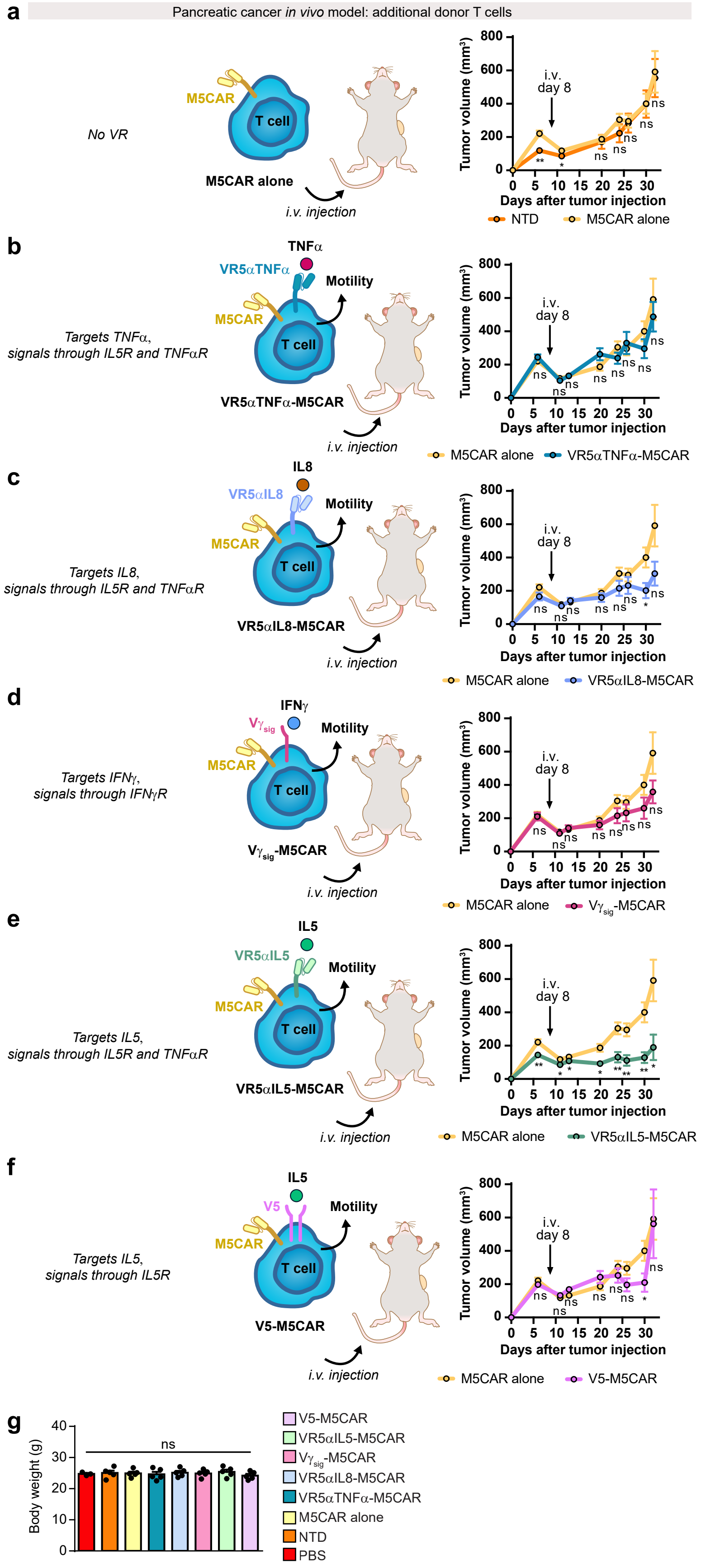

**Extended Data Fig. 6 | a,** $2x10^6$ ASPC1 pancreatic cancer tumors pre-mixed in 1:1 Matrigel:PBS were subcutaneously (s.c.) engrafted into 8-12 week old NSG mice. Tumor volumes were measured twice a week using digital calipers. When tumors were palpable (100 – 250 $mm^3$), mice were randomized and blindly treated (8 days later) with a single intravenous (i.v.) dose of $3x10^6$ NTD or M5CAR T cells. Tumor volumes were calculated as $(L \times W^2) \times 0.5$ plotted as $mm^3$. **b,** ASPC1-bearing mice were treated with M5CAR alone or VR5αTNFα-M5CAR T cells (as in panel **a**). **c,** ASPC1-bearing mice were treated with M5CAR alone or VR5αIL8-M5CAR T cells (as in panel **a**). **d,** ASPC1-bearing mice were treated with M5CAR alone or $V\gamma_{sig}$-M5CAR T cells (as in panel **a**). **e,** ASPC1-bearing mice were treated with M5CAR alone or VR5αIL5-M5CAR T cells (as in panel **a**). **f,** ASPC1-bearing mice were treated with M5CAR alone or V5-M5CAR T cells (as in panel **a**). **g,** Mice were weighed at the end of the study. For all figure panels, tumor volumes (**a-f**) and body weights (**g**) are plotted as mean ± SEM ($n = 5$ mice per group, except for PBS for which $n = 3$). For panels (**a-f**), two-tailed student's $t$ test was used for statistical analysis (ns = not significant, $*P < 0.05$, $**P < 0.01$). For panel (**g**), ordinary one-way ANOVA with Tukey's multiple comparison test was used for statistical analysis (ns = not significant).

**Extended Data Fig. 7**

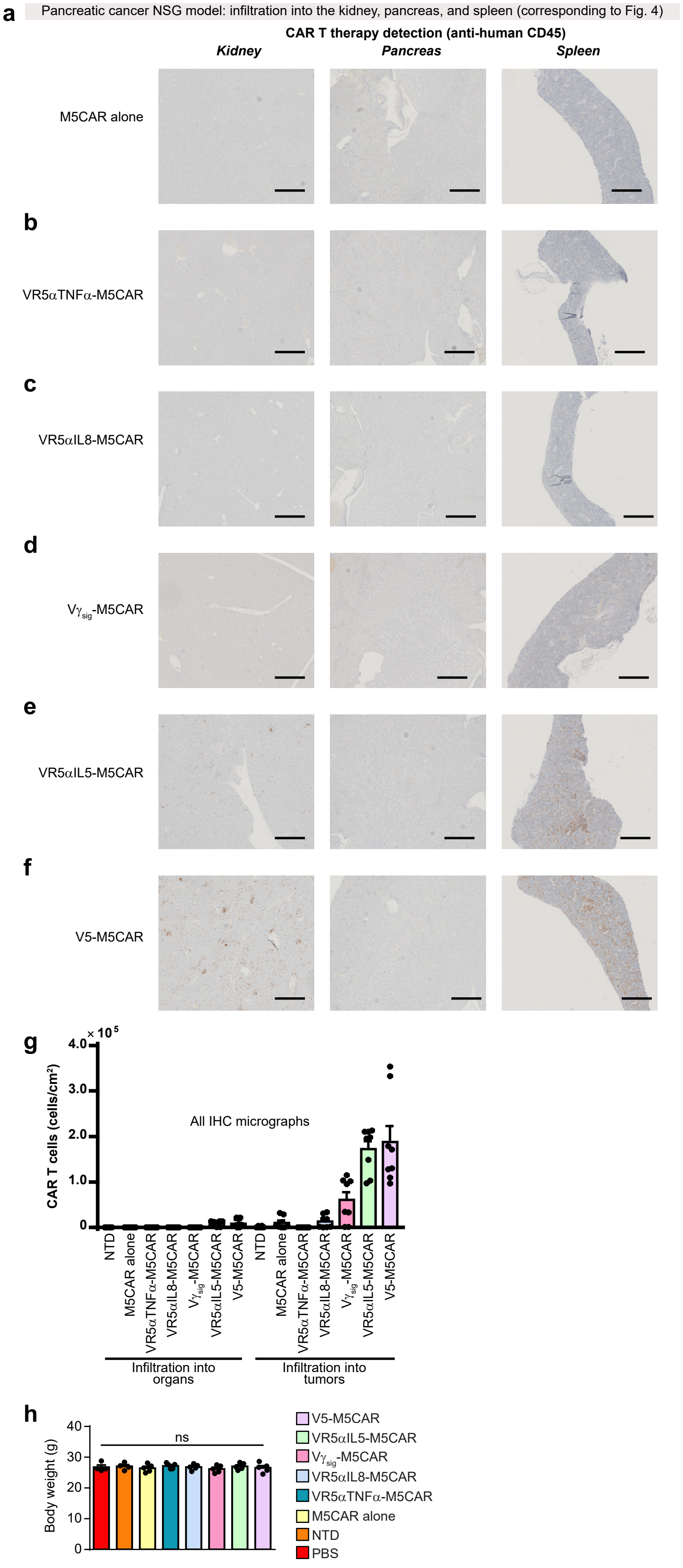

**Extended Data Fig. 7 | a,** ASPC1 pancreatic tumors were subcutaneously engrafted into NSG mice and measured as described in Fig. 4. Mice bearing ASPC1 tumors were treated with M5CAR T cells as described in Fig. 4. The kidney, pancreas, and spleen were harvested from day-25 mice. Organs were sectioned and IHC-stained with anti-human CD45 to detect T cell therapies as in Fig. 4. **b,** Mice bearing ASPC1 tumors were treated with VR5α TNFα-M5CAR T cells as described in Fig. 4. The kidney, pancreas, and spleen were sectioned and IHC-stained with anti-human CD45 to detect T cell therapies as in Fig. 4. **c,** ASPC1-bearing mice were treated with VR5α IL8-M5CAR T cells as described in Fig. 4. The kidney, pancreas, and spleen were sectioned and IHC-stained with anti-human CD45 for CAR detection. **d,** ASPC1-bearing mice were treated with $V\gamma_{sig}$-M5CAR T cells as described in Fig. 4. The kidney, pancreas, and spleen were sectioned and IHC-stained with anti-human CD45. **e,** ASPC1-bearing mice were treated with VR5α IL5-M5CAR T cells as described in Fig. 4. The kidney, pancreas, and spleen were sectioned and IHC-stained with anti-human CD45 to detect human T cells as in Fig. 4. **f,** ASPC1-bearing mice were treated with V5-M5CAR T cells as described in Fig. 4. The kidney, pancreas, and spleen were sectioned and IHC-stained with anti-human CD45 to detect T cell therapies as in Fig. 4. For figure panels (**a-f**), scale bars in IHC micrographs are 500 μm. **g,** Human CD45+ cell numbers were obtained as described in materials and methods. Tissue area was computationally obtained. Organ and tumor infiltration of human CD45+ cells per area were calculated and plotted. For calculation of cells/$cm^2$ in organs, $n$ = 6 for NTD or n = 8 for M5CAR alone and VR-M5CARs, with 2 non-consecutive tissue slides stained per mouse and 3-4 mice per group. For calculation of cells/$cm^2$ in tumors, $n$ = 6 for NTD or n = 8 for VR-M5CARs, with 2 non-consecutive tissue slides stained per mouse and 3-4 mice per group; n = 7 for M5CAR alone, with 2 non-consecutive tissue slides stained for 3 mice and 1 tissue slide stained for 1 mouse. **h,** Mouse body weights were collected at the end of the study. Body weights are plotted as mean ± SEM ($n$ = 5 mice per group, except for PBS and NTD for which $n$ = 4). Ordinary one-way ANOVA with Tukey's multiple comparison test was used for statistical analysis (ns = not significant).

**Extended Data Fig. 8**

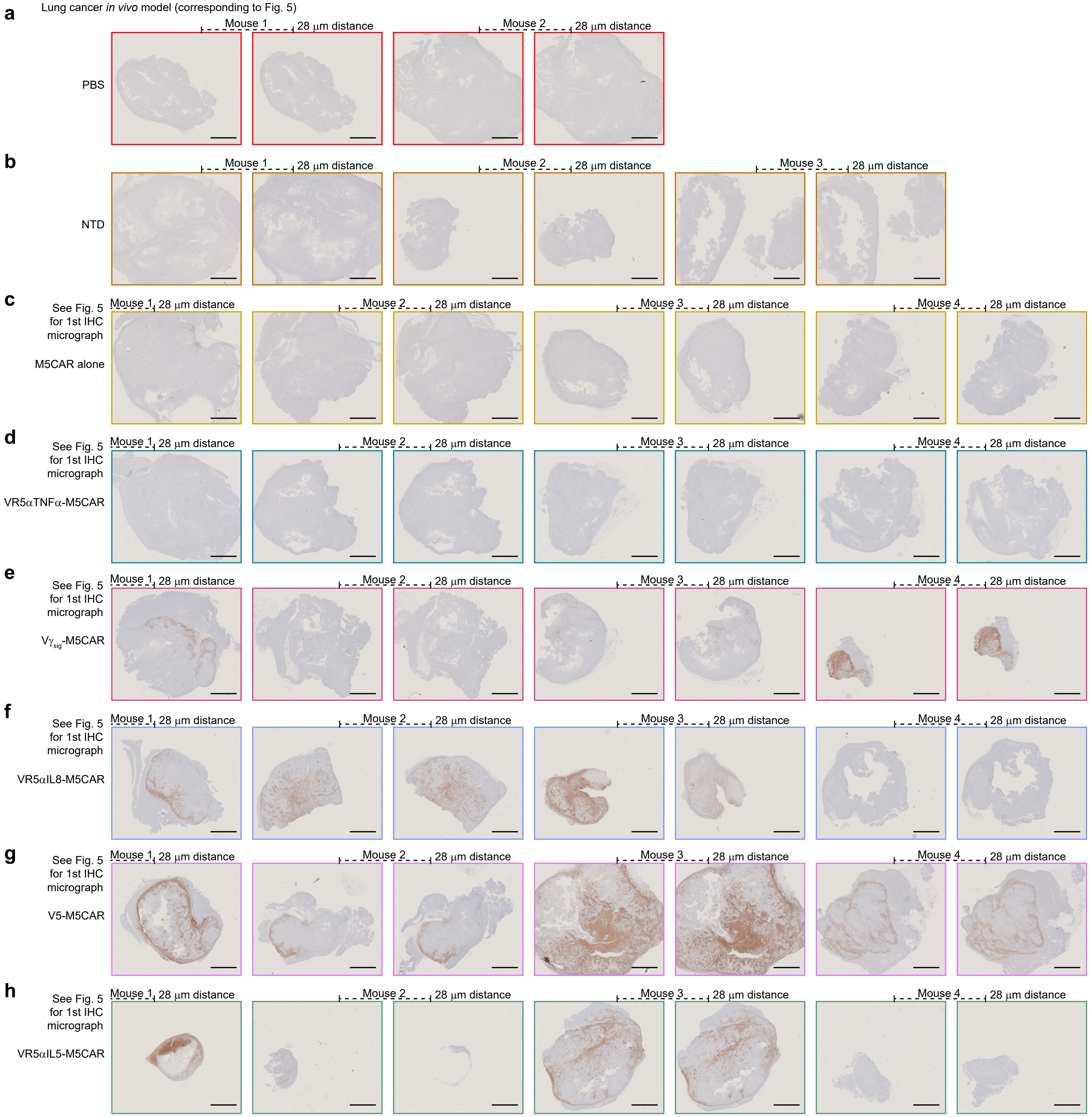

**Extended Data Fig. 8 | a-h**, CD45+ IHC micrographs of tumors from H226-bearing mice (corresponding to Fig. 5) treated with PBS (**a**), NTD T cells (**b**), M5CAR alone T cells (**c**), VR5αTNFα-M5CAR T cells (**d**), $V\gamma_{sig}$-M5CAR T cells (**e**), VR5αIL8-M5CAR T cells (**f**), V5-M5CAR T cells (**g**), or VR5αIL5-M5CAR T cells (**h**). Scale bars in IHC micrographs are 2 mm.

**Extended Data Fig. 9**

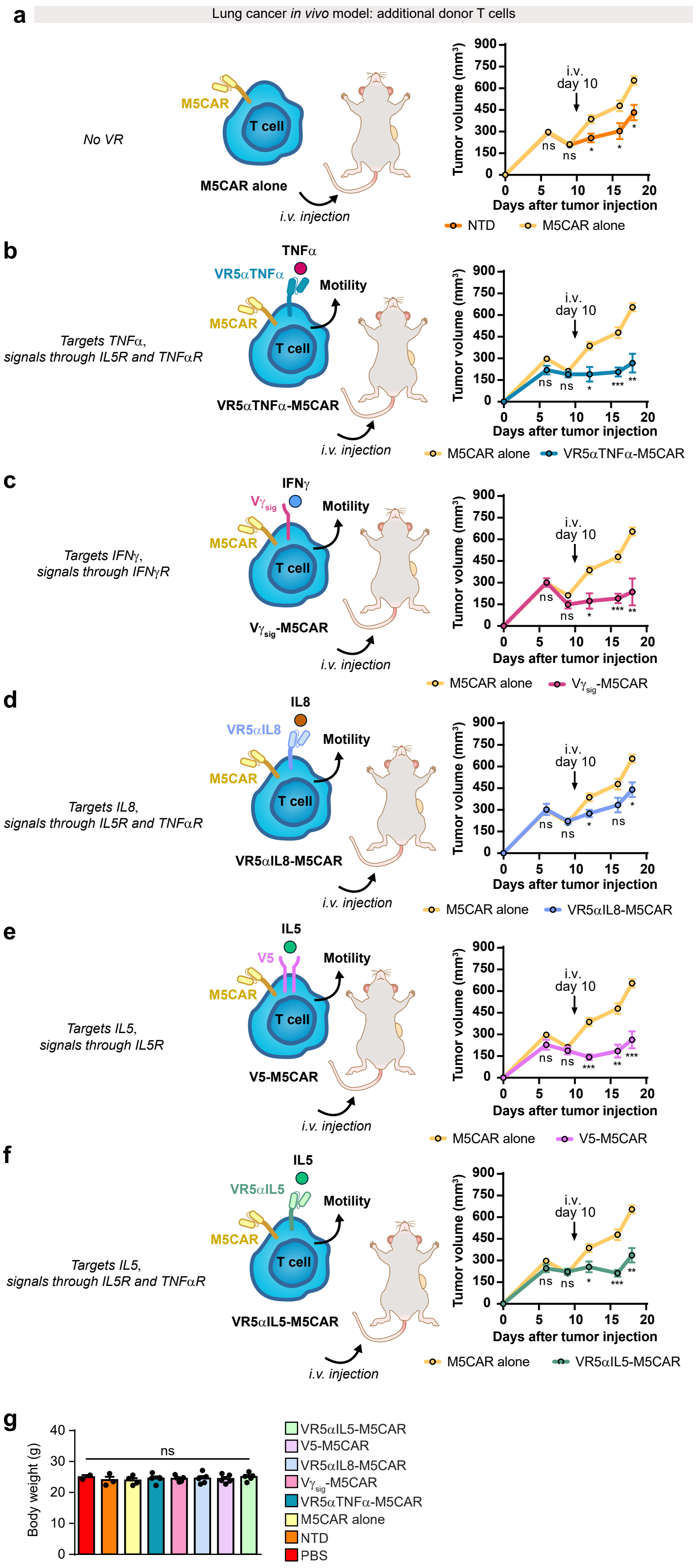

**Extended Data Fig. 9 | a,** 5x$10^6$ H226 lung cancer tumors pre-mixed in 1:1 Matrigel:PBS were engrafted s.c. into 8-12 week old NSG mice. Tumor volumes were measured twice a week using digital calipers. When tumors were palpable (100 – 250 $mm^3$), mice were randomized and blindly treated (10 days later) with a single i.v. dose of 3x$10^6$ NTD or M5CAR T cells. Tumor volumes were calculated as (L x $W^2$) × 0.5 plotted as $mm^3$. **b,** H226-bearing mice were treated with M5CAR alone or VR5αTNFα-M5CAR T cells (as in panel **a**). **c,** Mice bearing H226 tumors were treated with M5CAR alone or $V\gamma_{sig}$-M5CAR T cells (as in panel **a**). **d,** Mice bearing H226 tumors were treated with M5CAR alone or VR5αIL8-M5CAR T cells (as in panel **a**). **e,** H226-bearing mice were treated with M5CAR alone or V5-M5CAR T cells (as in panel **a**). **f,** Mice bearing H226 tumors were treated with M5CAR alone or VR5αIL5-M5CAR T cells (as in panel **a**). **g,** Mouse body weights were collected at the end of the study. For all figure panels, tumor volumes (**a-f**) and body weights (**g**) are plotted as mean ± SEM ($n$ = 5 mice per group, except for PBS for which $n$ = 2, NTD for which $n$ = 3, and M5CAR alone and VR5αTNFα-M5CAR for which $n$ = 4). For panels (**a-f**), two-tailed student's $t$ test was used for statistical analysis (ns = not significant, $*P < 0.05$, $**P < 0.01$). For panel (**g**), ordinary one-way ANOVA with Tukey's multiple comparison test was used for statistical analysis (ns = not significant).

**Extended Data Fig. 10**

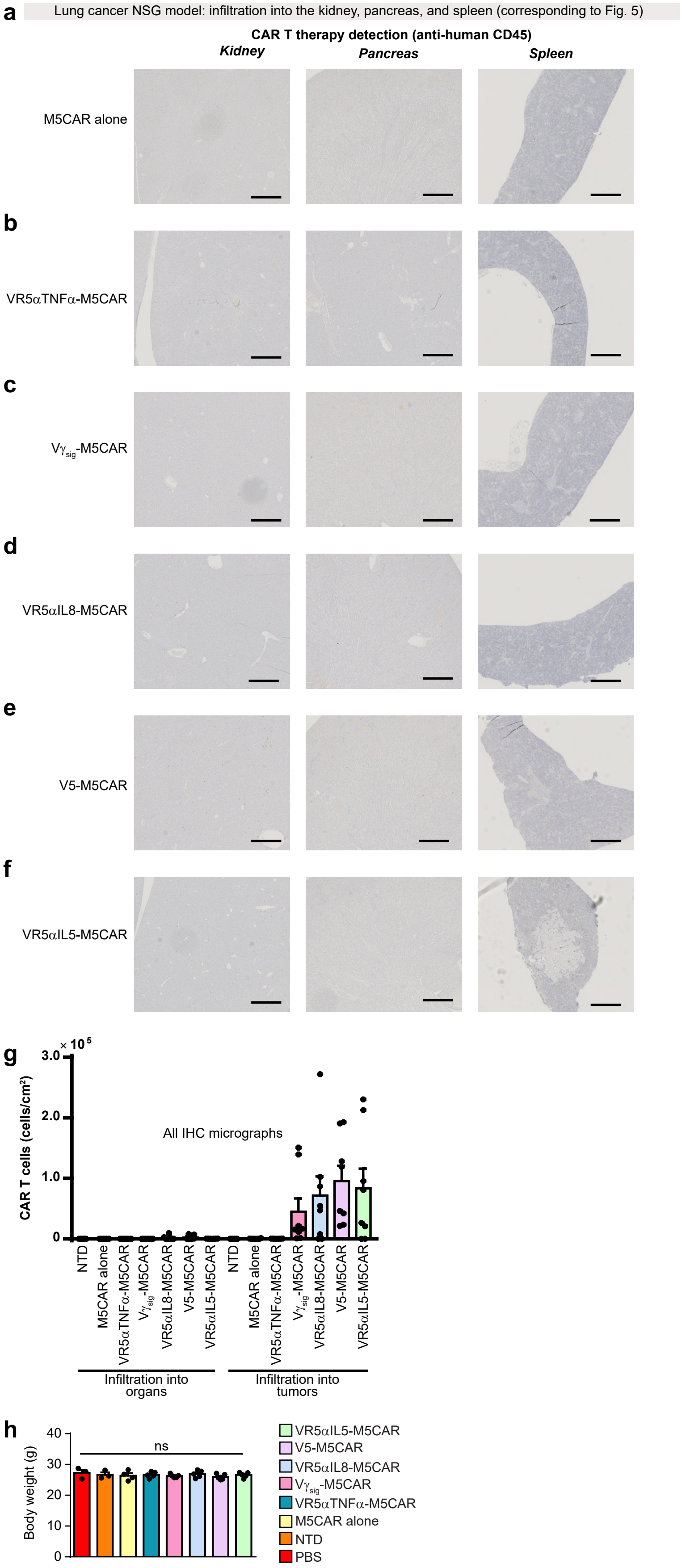

**Extended Data Fig. 10 | a,** H226 lung tumors were subcutaneously engrafted into NSG mice and measured as described in Fig. 5. Mice bearing H226 tumors were treated with M5CAR T cells as described in Fig. 5. The kidney, pancreas, and spleen were harvested from day-19 mice. Organs were sectioned and IHC-stained with anti-human CD45 to detect T cell therapies as in Fig. 5. **b,** Mice bearing H226 tumors were treated with VR5α TNFα-M5CAR T cells as described in Fig. 5. The kidney, pancreas, and spleen were sectioned and IHC-stained with anti-human CD45 to detect T cell therapies as in Fig. 5. **c,** H226-bearing mice were treated with $V\gamma_{sig}$-M5CAR T cells as described in Fig. 5. The kidney, pancreas, and spleen were sectioned and IHC-stained with anti-human CD45 for CAR detection. **d,** H226-bearing mice were treated with VR5α IL8-M5CAR T cells as described in Fig. 5. The kidney, pancreas, and spleen were sectioned and IHC-stained with anti-human CD45. **e,** H226-bearing mice were treated with V5-M5CAR T cells as described in Fig. 5. The kidney, pancreas, and spleen were sectioned and IHC-stained with anti-human CD45 to detect T cell therapies as in Fig. 5. **f,** Mice bearing H226 tumors were treated with VR5α IL5-M5CAR T cells as described in Fig. 5. The kidney, pancreas, and spleen were sectioned and IHC-stained with anti-human CD45 to detect T cell therapies as in Fig. 5. For figure panels (**a-f**), scale bars in IHC micrographs are 500 μm. **g,** Human CD45+ cell numbers were obtained as described in materials and methods. Tissue area was computationally obtained. Organ and tumor infiltration of human CD45+ cells per area were calculated and plotted. For calculation of cells/$cm^2$ in organs, $n$ = 6 for NTD or n = 8 for VR-M5CARs, with 2 non-consecutive tissue slides stained per mouse and 3-4 mice per group; n = 7 for M5CAR alone, with 2 non-consecutive tissue slides stained for 3 mice and 1 tissue slide stained for 1 mouse. For calculation of cells/$cm^2$ in tumors, $n$ = 6 for NTD or n = 8 for M5CAR alone and VR-M5CARs, with 2 non-consecutive tissue slides stained per mouse and 3-4 mice per group. **h,** Mouse body weights were collected at the end of the study. Body weights are plotted as mean ± SEM ($n$ = 5 mice per group, except for PBS for which $n$ = 2, NTD for which $n$ = 3, and M5CAR alone for which $n$ = 4). Ordinary one-way ANOVA with Tukey's multiple comparison test was used for statistical analysis (ns = not significant).
